# Stable centromere association of the yeast histone variant Cse4 requires its essential N-terminal domain

**DOI:** 10.1101/2024.07.24.604937

**Authors:** Andrew R. Popchock, Sabrine Hedouin, Yizi Mao, Charles L. Asbury, Andrew B Stergachis, Sue Biggins

## Abstract

Chromosome segregation relies on kinetochores that assemble on specialized centromeric chromatin containing a histone H3 variant. In budding yeast, a single centromeric nucleosome containing Cse4 assembles at a sequence-defined 125 bp centromere. Yeast centromeric sequences are poor templates for nucleosome formation *in vitro*, suggesting the existence of mechanisms that specifically stabilize Cse4 nucleosomes *in vivo*. The extended Cse4 N-terminal tail binds to the chaperone Scm3, and a short essential region called END within the N-terminal tail binds the inner kinetochore complex OA. To address the roles of these interactions, we utilized single molecule fluorescence assays to monitor Cse4 during kinetochore assembly. We found that OA and Scm3 independently stabilize Cse4 at centromeres via their END interaction. Scm3 binding to the Cse4 END is enhanced by Ipl1/Aurora B phosphorylation, identifying a previously unknown role for Ipl1 in ensuring Cse4 stability. Strikingly, an Ipl1 phosphomimetic mutation in the Cse4 END enhances Scm3 binding and can restore Cse4 recruitment in mutants defective in OA binding. Together, these data suggest that a key function of the essential Cse4 N-terminus is to ensure Cse4 localization at centromeres.

## INTRODUCTION

During cell division, chromosomes are replicated to form sister chromatids and then segregated to daughter cells. Chromosome segregation harnesses the forces generated by spindle microtubules through the kinetochore, a conserved megadalton protein machine. Kinetochores contain dozens of multiprotein subunits that assemble on centromeric chromatin containing a specialized histone H3 variant called CENP-A [1–3]. Remarkably, kinetochores must assemble *de novo* at centromeres every cell cycle after replication [4–6]. Most eukaryotes have large regional centromeric regions containing interspersed CENP-A and canonical H3 nucleosomes that serve as the platform for the recruitment of chromosome-adjacent inner kinetochore proteins to form the <u>c</u>onstitutive <u>c</u>entromere <u>a</u>ssociation <u>n</u>etwork (CCAN) [7–9], which then facilitates assembly of the outer kinetochore that interacts with spindle microtubules [10, 11]. In contrast, budding yeast have a unique point centromere that is sequence-defined and recruits a single CENP-A nucleosome that mediates assembly of the entire kinetochore [12–14]. Because yeast kinetochores assemble on a short, defined sequence with a single nucleosome, they are an ideal model system to study kinetochores since key aspects of their architecture and function are conserved.

Although a single yeast centromeric nucleosome is sufficient for kinetochore assembly, yeast centromeric DNA is a poor template for the reconstitution of CENP-A (Cse4) nucleosomes using purified components *in vitro* [12, 15–17]. Reconstitution of yeast centromeric nucleosomes requires additional stabilizing factors to form a stable nucleosome-core particle [18], or alterations of the native sequence in more complete kinetochore reconstitutions [19, 20]. The inherent unfavourability for stable centromeric nucleosome formation is counter to their critical function of kinetochore assembly in cells but has been proposed as a possible regulatory function to exclude canonical nucleosomes [21]. Because centromere nucleosome assembly must occur rapidly and with high fidelity after replication, factors must exist that promote this process *in vivo*. Consistent with this requirement, we recently found that Cse4 stabilization requires two additional kinetochore proteins, the conserved histone chaperone Scm3 (HJURP) and the essential kinetochore complex OA (CENP-QU) [22], although the mechanisms by which they stabilize the centromeric nucleosome are not known.

Scm3 and the OA complex directly interact with Cse4. The Scm3 chaperone was initially found to bind to a Cse4 region in the histone fold domain containing residues 166-201 called the <u>c</u>entromere <u>a</u>ssociated targeting <u>d</u>omain (CATD) [23–26]. However, a recent study reported that Scm3 also binds to Cse4’s highly disordered extended N-terminal domain (NTD, Cse4-1-133) which contains the initial 133 residues of Cse4 [27]. Although the NTD is essential, it is absent from reconstituted structural models due to its intrinsic disorder, so the interactions between OA and Scm3 and the Cse4-NTD within the reconstituted inner kinetochore are unknown. The NTD contains a short essential region spanning residues 28-60, named the <u>e</u>ssential <u>N</u>-terminal <u>d</u>omain (END, Cse4-28-60) [28] and it is the target of several different types of PTMs [29–31], but the functional consequences of these modifications remain largely unknown. The END contains an essential OA binding interface [28, 30, 32, 33] and was also shown to undergo significant structural rearrangement in reconstitutions containing Cse4-NTD bound to Scm3 [27]. The additional Scm3 interaction site in Cse4 expands the potential functional roles of Scm3 at the kinetochore beyond its canonical role in Cse4 centromere targeting through the conserved CATD domain. This additional function may depend upon its interaction with the N-terminal tail of Cse4, a region which has been shown to play a role in the localization of CENP-A to centromeres in *C. elegans* [34, 35]. This is consistent with our recent findings that Scm3 can stabilize the interaction of Cse4 with centromeric DNA after initial association during *de novo* kinetochore assembly by an unknown mechanism [22]. An additional function for Scm3 at centromeres may also explain its behavior in cells, where it has been shown to be in constant exchange at kinetochores throughout mitosis, long after centromeric targeting and stable Cse4 incorporation [4].

To elucidate the function of Scm3 and OA binding to the essential N-terminus of Cse4, we utilized our recently developed single molecule fluorescence assay that assembles centromeric nucleosomes in yeast lysates. In contrast to conventional biochemical reconstitutions that do not achieve stable centromeric nucleosomes without alterations or stabilizing techniques, Cse4 is rapidly assembled onto centromeric DNA sequences in yeast extract and these native nucleosomes are remarkably stable when removed from extract [22]. Here we show that the END region is required for stable Cse4 association with centromeric DNA and requires the independent recruitment of both OA and Scm3. In cells, we find that disruption of END binding to OA causes a reduction of Cse4 centromeric levels that can be rescued via enhancing the END binding of Scm3. Taken together, our data indicate that the Cse4 END domain not only recruits kinetochore proteins, but that END recruitment of these proteins stabilizes the centromeric nucleosome. We propose that this multistep stabilization mechanism not only ensures that cells assemble a single Cse4 nucleosome at centromeres, but also makes it difficult to stabilize ectopic centromeric nucleosomes that could lead to genomic instability.

## RESULTS

### OA contributes to Cse4 stability in cells

To test if the recruitment of OA contributes to Cse4 stability in cells, we utilized a technique called Fiber-seq that enables nucleosome mapping at single-molecule level on chromatin fibers [36, 37]. Briefly, spheroplasted cells were permeabilized and treated with a nonspecific N6-adenine DNA methyltransferase (m6A-MTase) that methylates accessible (non-protein bound) DNA (**Figure 1A**). Individual m6A-stenciled chromatin fibers are sequenced using long-read single-molecule sequencing to >100x genomic coverage to identify single-molecule protein occupancy events at single-molecule and near single-nucleotide resolution (**Figure 1A**). The entirety of each point centromere in budding yeast is readily captured along a sequenced read, which are >10kb in length, and as this is a genome-wide method, each of the 16 centromeres was thoroughly captured by sequenced fibers (∼1100 fibers for WT cells and ∼500 fibers for mutants). Consequently, this approach enabled the direct quantification of the average steady-state absolute occupancy of the centromeric nucleosome across individual fibers in yeast cells. We began by testing whether differences in Cse4 occupancy levels at centromeres could be detected under the most stringent conditions, when Cse4 is depleted using a *cse4-AID* system. We performed Fiber-seq on WT and Cse4-depleted cells and found nearly full nucleosome occupancy (92.2 %) at centromeres in WT cells, which was markedly reduced to 41.6 % in *cse4-AID* cells (**Figure 1B**). We then depleted OA in cells using an *okp1-AID* strain to determine if Cse4 nucleosome occupancy was affected. After OA depletion, nucleosome occupancy levels at centromeres were significantly reduced (56.0 %) to levels comparable to those after Cse4 depletion (**Figure 1B**). This indicated a loss of Cse4 at centromeres after OA depletion that was consistent with our previous observations in *de novo* reconstitutions [22], suggesting a role for OA in retention of Cse4 at centromeres.

**Figure 1.**
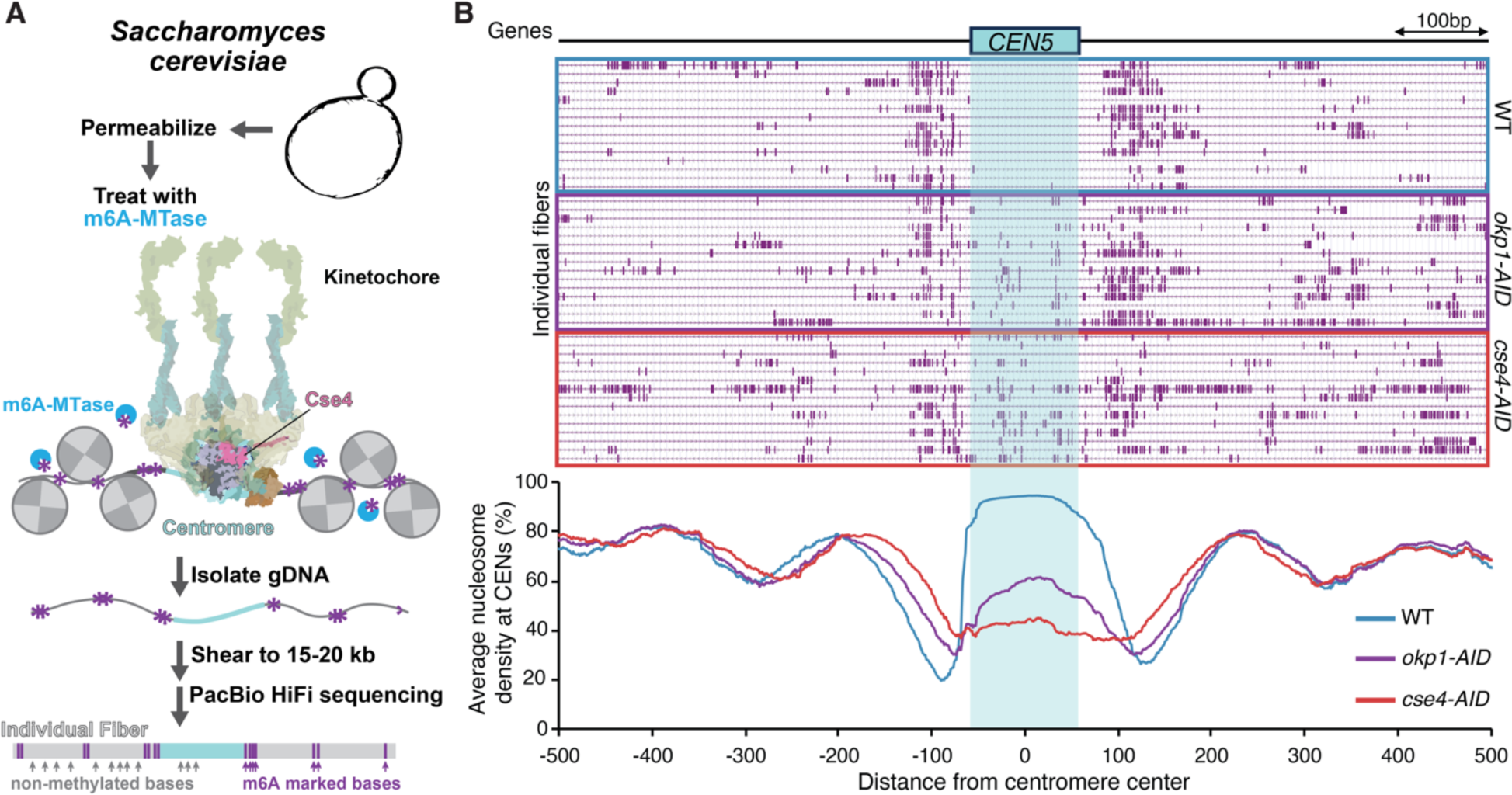
Depletion of Okp1 significantly reduces Cse4 nucleosome density at centromeres in cells. A. Schematic representing experimental design for Fiber-seq of *Saccharomyces cerevisiae* including example centromeric loci including the kinetochore-bound centromeric nucleosome. B. Example Fiber-seq reads of chromatin fibers with methylated bases shown in purple through centromere regions of chromosome V in WT (SBY3, top), *okp1-AID* (SBY15124, middle) and *cse4-AID* (SBY22656, bottom) cells. Bottom graph indicates calculated average nucleosome density for all chromosomes centered at centromeres from Fiber-seq genomic DNA analysis from *CSE4-GFP* (blue), *okp1-AID* (purple) and *cse4-AID* (red) cells.

### The Cse4 END is critical for Cse4 recruitment to centromeric DNA

Because OA is required for stable Cse4 association with centromeric DNA *in vivo*, we asked whether this is mediated by its essential interaction with the END [32]. To test this, we first asked whether the END contributes to Cse4 stability at the centromere. We monitored Cse4 centromeric recruitment using a recently developed single molecule TIRFm *de novo* kinetochore assembly assay. Briefly, fluorescently labeled centromeric DNA templates (CEN DNA) are sparsely attached to a coverslip surface and then cell extract containing endogenously tagged fluorescent kinetochore protein(s) is introduced into a flow chamber and incubated for 90 min (**Figure 2A**). After incubation to allow kinetochore assembly, the extract is washed away and the colocalization of the target fluorescent protein with the labeled CEN DNA is quantified (**Figure 2A**). We introduced an ectopic copy of GFP-tagged wild type Cse4 (*pGAL*-*CSE4-GFP)*, or a deletion mutant lacking the END, Cse4^ϕλEND^ (*pGAL-cse4^ϕλEND^-GFP*) (**Figure 2B**), under a galactose inducible promoter to overexpress them during a mitotic arrest. Ectopic expression was needed because the Cse4 END is essential for viability. The proteins were overexpressed to similar levels (**Supplemental Figure 1A**). After incubation with *pGAL-CSE4-GFP* extract for 90’, ∼50 % of the CEN DNA exhibited co-localization with overexpressed Cse4-GFP, similar to endogenous Cse4-GFP co-localization [22]. In contrast, the endpoint colocalization of overexpressed Cse4^ΔEND^-GFP was significantly reduced to 12 % (**Figure 2B**), indicating a role for the END in stable Cse4 centromere recruitment. Because prior structural studies with reconstituted nucleosomes do not provide a mechanism for how the END would stabilize Cse4 and also do not use native centromere sequences [19, 20], we asked whether the native kinetochore assembly pathway differs from what occurs during conventional reconstitutions. To do this, we tested the non-native centromere sequence used in the most recent and complete structural studies (C0N3 DNA) in TIRFM kinetochore assembly assays (**Figure 2C**) [20]. We found that despite slightly enhanced recruitment of the CBF3 component Ndc10 (**Supplemental Figure 1B**), Cse4 levels were significantly lower when compared to the native centromere sequence (CEN DNA, **Figure 2C**), suggesting that the assembly pathway of reconstituted kinetochores using purified proteins may differ from those containing native components. This significant drop in Cse4 levels observed on C0N3 DNA is much greater than variance we would expect between native centromere sequences based on previous observations and is comparable to mutant centromere sequences that are unstable in cells [22]. Together, these data identify a critical role for the Cse4 END domain in its centromere association, a finding that was not previously apparent from structural reconstitutions.

**Figure 2.**
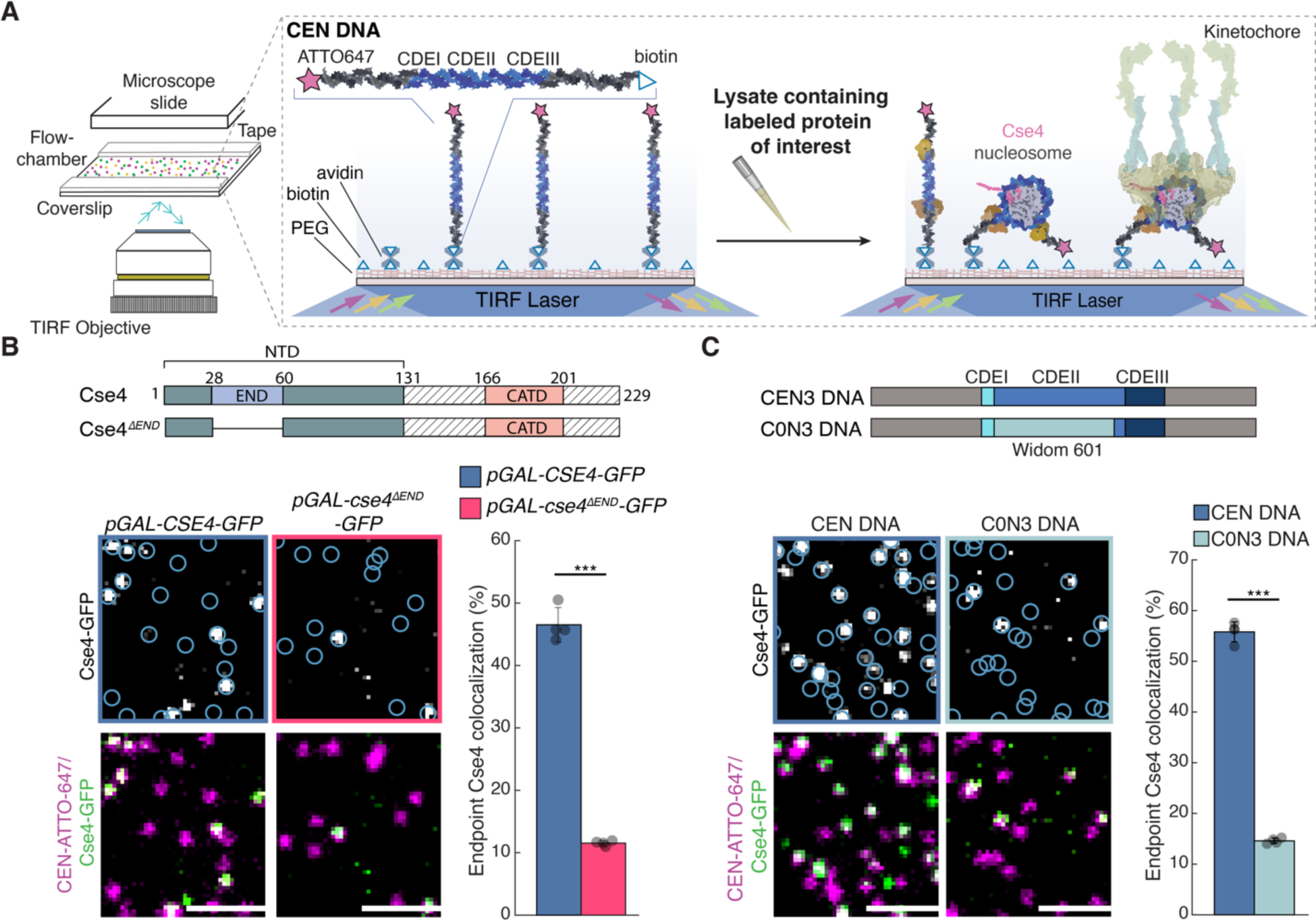
Cse4 centromeric localization depends on both the Cse4 END region and native centromeric sequence. A. Schematic diagram of *de novo* TIRFm kinetochore assembly assays (adapted from [22]). B. Cse4 construct schematic (top) and example images of TIRFM endpoint colocalization assays. Top panels show visualized Cse4-GFP on CEN DNA in *pGAL-CSE4-GFP* (SBY22273) extracts (top-left) or *pGAL-cse4^τιEND^-GFP* (SBY22803) extracts (top-right) with colocalization shown in relation to identified CEN DNA in blue circles. Bottom panels show respective overlays of CEN DNA channel (magenta) with Cse4-GFP (green). Scale bars 2 μm. Graph indicates Cse4-GFP endpoint colocalization with CEN DNA in extracts from *pGAL-CSE4-GFP* or *pGAL-cse4-τ<END-GFP* genetic backgrounds (47 ± 2.8%, 12 ± 0.4%, avg ± s.d. n = 4 experiments, each examining ∼ 1,000 DNA molecules from different extracts, *** indicates significant difference with two-tailed *P*-value of 1.4E-4). C. DNA template schematic (top) and example images of TIRFM endpoint colocalization assays. Top panels show visualized Cse4-GFP on CEN DNA (top-left panel), or on C0N3 DNA (top-right panel) in *CSE4-GFP* (SBY21863) extracts with colocalization shown in relation to identified DNAs in blue circles. Bottom panels show overlay of CEN or C0N3 DNA channel (magenta) with Cse4-GFP (green), Scale bars 2 μm. Graph indicates quantification of Cse4-GFP endpoint colocalization with CEN DNA or C0N3 DNA (56 ± 2.0%, 15 ± 0.5%, avg ± s.d. n = 4 experiments, each examining ∼ 1,000 DNA molecules from different extracts, *** indicates significant difference with two-tailed *P*-value of 3.5E-5).

### OA binding to Cse4 END stabilizes the centromeric nucleosome

To determine whether Cse4 stabilization by the END domain is due to its interaction with OA, we turned to recently reported Cse4 mutations at L41 that disrupt OA binding [32]. Consistent with that publication, we found that Cse4-NTD^L41A^ and Cse4-NTD^L41D^ disrupted OA binding in pull down assays (**Figure 3A**). Although the Cse4-L41D mutant more strongly disrupted OA binding, it was not used for further study because it is inviable [32] and the use of endogenous Cse4 alleles was necessary for monitoring Cse4 behavior with respect to other kinetochore proteins. We next analyzed OA localization to centromeres by performing bulk kinetochore assembly assays, where CEN DNA templates are linked to magnetic beads and directly incubated in cell extracts [14]. We found that cell extracts containing the Cse4^L41A^ mutant completely abolished OA complex recruitment (**Supplemental Figure 2A**), indicating that OA kinetochore assembly requires binding to the Cse4 END. Consistent with an additional role for OA in stabilizing Cse4, we found that Cse4 centromere recruitment was also significantly reduced in Cse4^L41A^ mutant extracts (**Supplemental Figure 2A**). However, the recruitment of Ndc10, which is necessary for Cse4 deposition, was not affected (**Supplemental Figure 2A**). To quantify the Cse4 recruitment defect, we performed TIRFM endpoint colocalization assays. Consistent with the bulk assembly assays, Cse4^L41A^-GFP endpoint colocalization was significantly lower than Cse4-GFP (7 % vs 55 % respectively, **Figure 3B**). To confirm this was due to disruption of the END:OA binding interface (**Supplemental Figure 2B**), we introduced an orthogonal mutation in the OA complex component protein Ame1 at residue I195 (*ame1^I195Y^*), which was previously shown to abrogate OA binding to the Cse4 END at the same interface as the Cse4 L41 mutants via disruption in hydrophobic packing [32]. In endpoint colocalization assays, Cse4-GFP levels were significantly reduced in extracts containing the Ame1^I195Y^ mutant (17 % vs 55 % respectively, **Figure 3B**). Together, these results indicate that END recruitment of OA is needed for stable Cse4 association with centromeric DNA.

**Figure 3.**
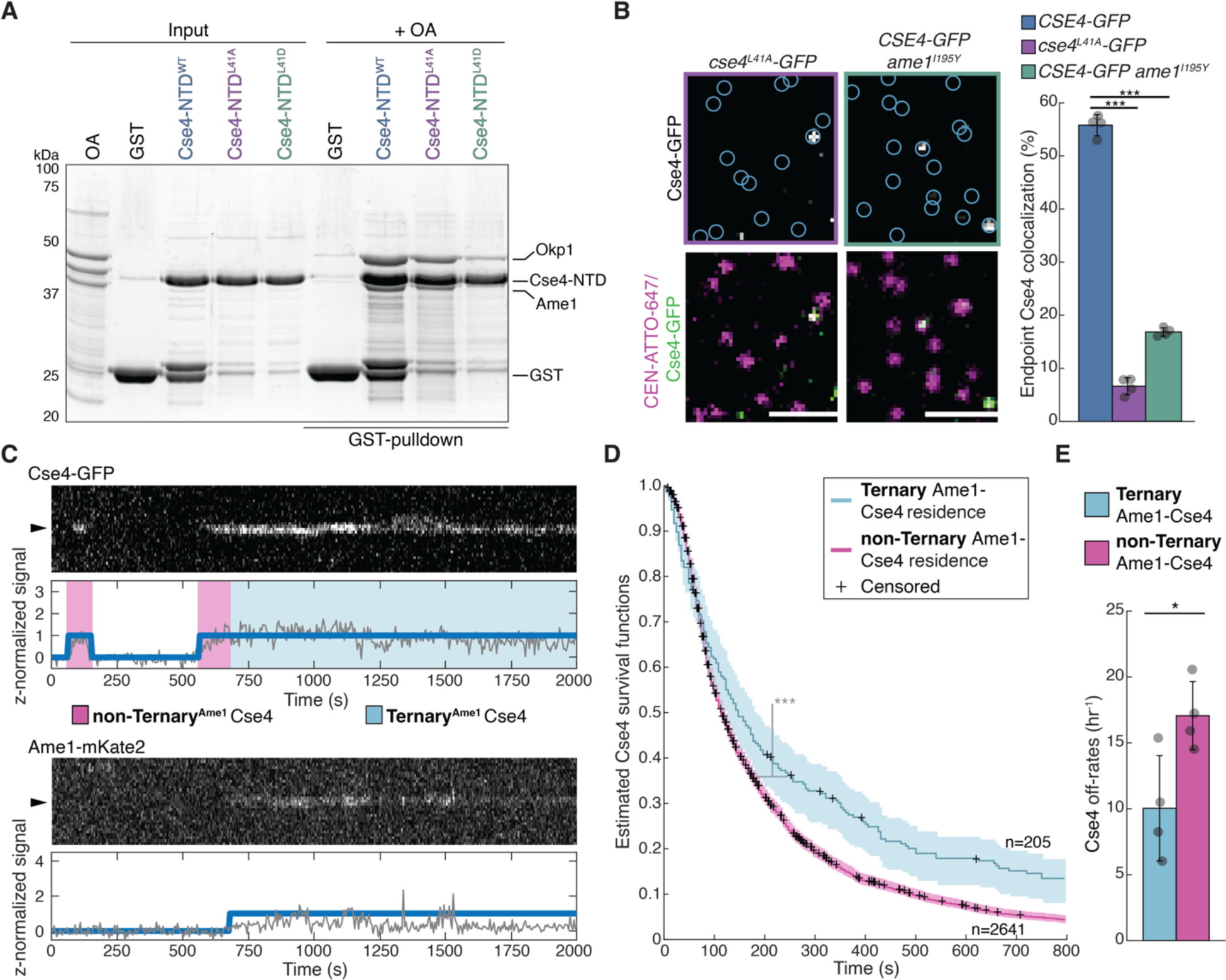
OA recruitment by Cse4 END is required for stable Cse4 localization to centromeric DNA. A. SDS-PAGE of GST pulldown assays of immobilized Cse4-NTD^WT^, Cse4-NTD^L41A^ and Cse4-NTD^L41D^ to test binding of recombinant OA. B. Example images of TIRFM endpoint colocalization assays. Top panels show visualized Cse4-GFP on CEN DNA in *cse4^L41A^-GFP* (SBY22811) extracts (top-left panel) or *CSE4-GFP ame1^I195Y^* (SBY23105) extracts (top-right panel) with colocalization shown in relation to identified CEN DNA in blue circles. Bottom panels show overlay of CEN DNA channel (magenta) with Cse4 -GFP (green), scale bars 2 μm. Graph indicates quantification of Cse4-GFP endpoint colocalization with CEN DNA in extracts from *CSE4-GFP*, *cse4^L41A^-GFP* or *CSE4-GFP ame1^I195Y^* genetic backgrounds (56 ± 2.0 %, 6.6 ± 1.6 % and 16.8 ± 0.8 % avg ± s.d. n = 4 experiments, each examining ∼ 1,000 DNA molecules from different extracts, *** indicates significant difference with two-tailed *P*-value between *CSE4-GFP* and *cse4^L41A^-GFP* of 2.1E-8 and between *CSE4-GFP* and *CSE4-GFP ame1^I195Y^* of 3.5E-6). C. Representative residence lifetime assay trace of Cse4-GFP and Ame1-mKate2 on a single CEN DNA in *CSE4-GFP AME1-mKate2 extract* (SBY22244). Top panel includes kymograph of Cse4 (top-488 nm) in relation to single identified CEN DNA (arrow), with normalized intensity trace (gray-bottom) as well as identified residences (blue). Bottom panel includes kymograph of Ame1 (bottom-561 nm) in relation to the same identified CEN DNA (arrow), with normalized intensity trace (gray-bottom) as well as identified residences (blue). Images acquired every 5 s with normalized fluorescence intensity shown in arbitrary units. D. Estimated survival function plots of Kaplan–Meier analysis of the lifetimes of Ternary Ame1-Cse4 residences on CEN DNA (blue— median lifetime of 147 s, n = 205 over four experiments of ∼ 1,000 DNA molecules using different extracts) and non-Ternary Ame1-Cse4 residences on CEN DNA (purple—median lifetime of 113 s, n = 2641 over four experiments of ∼ 1,000 DNA molecules using different extracts). 95% confidence intervals indicated (dashed lines), right-censored lifetimes (plus icons) were included with equivalent weighting in survival function estimates, *** indicates significant difference two-tailed *P*-value of 3.0e-06 as determined by log-rank test. E. Graph indicates quantification of the estimated off-rates of Ternary Ame1-Cse4 and non-Ternary Ame1-Cse4 residences on CEN DNA (10.0 h^−1^ ± 4.0 h^−1^ and 17.1 ± 2.6 h^−1^ respectively, avg ± s.d. n = 3598 over four experiments of ∼ 1,000 DNA molecules, * indicates significant difference with two-tailed *P*-value of .03).

To further explore Cse4 stabilization by OA binding, we sought to dissect the behavior of Cse4 on centromeric DNA in relation to OA. To do this, we utilized our previously developed time-lapse assay that enables direct and simultaneous monitoring of two orthogonally labeled fluorescent kinetochore proteins in cell extract with CEN DNA over time [22]. Continuous monitoring allows us to analyze more complex and dynamic binding behavior based on simultaneous protein-protein colocalization. We therefore orthogonally labeled Ame1-mKate2 and Cse4-GFP and simultaneously monitored their behavior to identify instances when they formed a ternary complex on centromeric DNA (**Figure 3C**). We then compared the residence times of Cse4 on CEN DNA in the presence of OA (Ternary Ame1-Cse4) or absence of OA (non-Ternary Ame1-Cse4) (**Figure 3C**). Kaplan-Meir analysis revealed a significant increase in the median lifetime of Ternary Ame1-Cse4 versus non-Ternary Ame1-Cse4 (147 s vs. 113 s respectively, **Figure 3D**). Consistent with this, the off-rates of Ternary Ame1-Cse4 were significantly slower (indicating more stable association) on centromeric DNA when compared to non-Ternary Ame1-Cse4 (10.0 h^−1^ vs 17.1 h^−1^, **Figure 3E**). These data indicate that the formation of a Cse4-OA complex stabilizes Cse4 on CEN DNA, providing an explanation for the reduced endpoint localization of Cse4^L41A^ to CEN DNA.

### Conserved mitotic kinase Ipl1 contributes to stable Cse4 centromeric localization via END domain phosphorylation

The END domain is phosphorylated by the Aurora B (Ipl1) kinase on Cse4-S40 [29], so we next asked whether Ipl1 regulates Cse4 centromeric localization. To test this, we analyzed kinetochore assembly in bulk assays when extracts had been depleted of Ipl1 (*ipl1-AID*) and found Cse4 recruitment to centromeric DNA was reduced (**Figure 4A**). We confirmed this requirement for Ipl1 in Cse4 centromeric localization in TIRFM assays (**Figure 4B**). To test whether the Ipl1-dependent requirement was dependent upon Cse4 END phosphorylation, we asked if a phosphomimetic mutant could restore Cse4 localization in the absence of Ipl1. Strikingly, Cse4^S40D^ restored Cse4 levels in Ipl1 depleted extracts in both bulk assembly assays and TIRFM assembly (**Figure 4C, 4D**). Taken together, these results suggest that Ipl1 plays a key regulatory role in ensuring Cse4 is stably associated with centromeres through the END domain.

**Figure 4.**
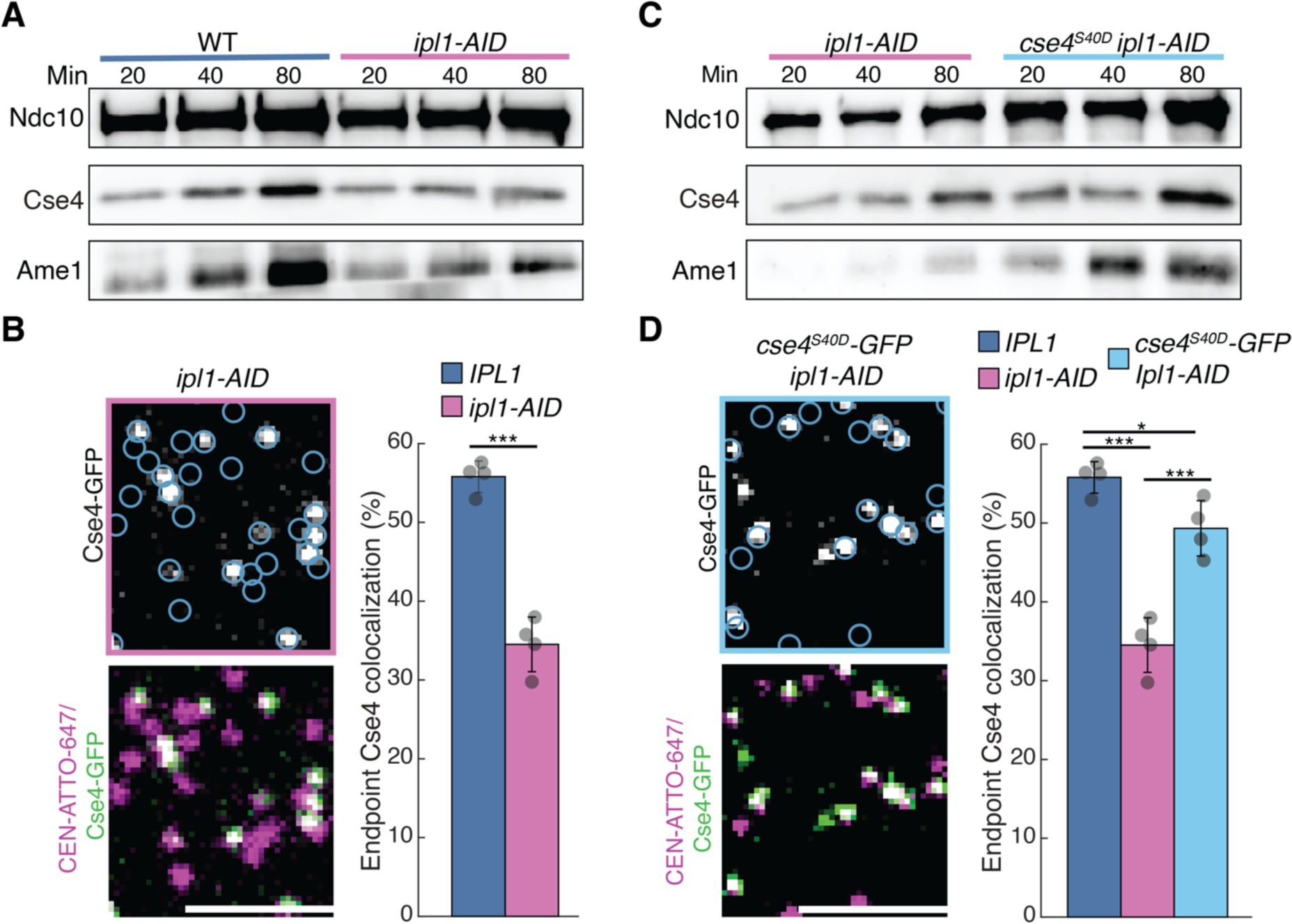
Ipl1 contributes to Cse4 localization via END phosphorylation. A. Immunoblots of bulk kinetochore assembly assays on centromeric DNA in WT (SBY4) cell extract (left) and *ipl1-AID* (SBY17186) extracts (right). DNA-bound proteins were analyzed by immunoblotting with the indicated antibodies. B. Example images of TIRFM endpoint colocalization assays. Top panels show visualized Cse4-GFP on CEN DNA *CSE4-GFP ipl1-AID* (SBY21972) extracts with colocalization shown in relation to identified CEN DNA in blue circles. Bottom panels show overlay of CEN DNA channel (magenta) with Cse4-GFP (green), Scale bars 3 μm. Graph indicates quantification of Cse4-GFP endpoint colocalization with CEN DNA in *IPL1* (SBY21863) or *ipl1-AID* extracts (56 ± 2.0%, 34 ± 3.5%, avg ± s.d. n = 4 experiments, each examining ∼ 1,000 DNA molecules from different extracts, *** indicates significant difference with two-tailed *P*-value of 1.3E-4). C. Immunoblots of bulk kinetochore assembly assays on centromeric DNA in *ipl1-AID* (SBY17186) extracts (left) and *cse4^S40D^ ipl1-AID* (SBY22431) extracts (right). D. Example images of TIRFM endpoint colocalization assays. Top panel shows visualized Cse4-GFP on CEN DNA in *cse4^S40D^-GFP ipl1-AID* (SBY20021) extracts with colocalization shown in relation to identified CEN DNA in blue circles. Bottom panel shows overlay of CEN DNA channel (magenta) with Cse4-GFP (green), Scale bars 3 μm. Graph indicates quantification of endpoint colocalization with CEN DNA of Cse4 in *cse4^S40D^-GFP ipl1-AID* extracts (light blue - 49 ± 3.5%, avg ± s.d. n = 4 experiments, each examining ∼ 1,000 DNA molecules from different extracts, *** indicates significant difference between *IPL1* and *ipl1-AID* with two-tailed *P*-value of 1.3E-4 and a difference between *ipl1-AID* and *cse4^S40D^-GFP ipl1-AID* with two-tailed *P*-value of 9.7E-4, * indicates significant difference between *IPl1* and *cse4^S40D^-GFP ipl1-AID* with two-tailed *P*-value of .02).

### Scm3 binds to the END domain and Ipl1 phosphorylation enhances the interaction

Because the Cse4 END domain binds to OA, we tested whether Ipl1 phosphorylation regulates OA association. To do this, we performed a pulldown assay with recombinant OA and were surprised to find that the Cse4-NTD^S40D^ mutant did not significantly alter OA binding (**Figure 5A**). However, it was recently reported that the Cse4 N-terminus also binds to the Scm3 chaperone [27], so we tested whether the END domain is required for Scm3 association in a pulldown assay. We generated recombinant GST-fusions to the entire Cse4 N-terminal domain (Cse4-NTD^WT^) as well as the NTD lacking the END region (Cse4-NTD^ΔEND^) to use as bait in pulldown assays (**Figure 5B**). We first confirmed that recombinant OA binding to the Cse4-NTD^WT^ was completely lost in the Cse4-NTD^ΔEND^ mutant, consistent with previous studies (**Figure 5B**). We then performed pulldown assays with recombinant Scm3 and found that it readily bound Cse4-NTD^WT^, but its binding was significantly disrupted in the Cse4-NTD^ΔEND^ mutant (**Figure 5B).** Because Cse4-S40 phosphorylation occurs near a unique acidic patch within the Cse4-NTD that was proposed to contribute to the specificity of the Scm3:Cse4-NTD interaction [27] and is somewhat distal to the hydrophobically packed Leu41 (**Supplemental Figure 2B**), we tested if the phosphomimetic Cse4-NTD^S40D^ enhanced Scm3:END binding in pulldown assays. Scm3 binding to Cse4-NTD^S40D^ was stimulated relative to Cse4-NTD under varying concentrations of purified recombinant Scm3 (**Figure 5C, D**).

**Figure 5.**
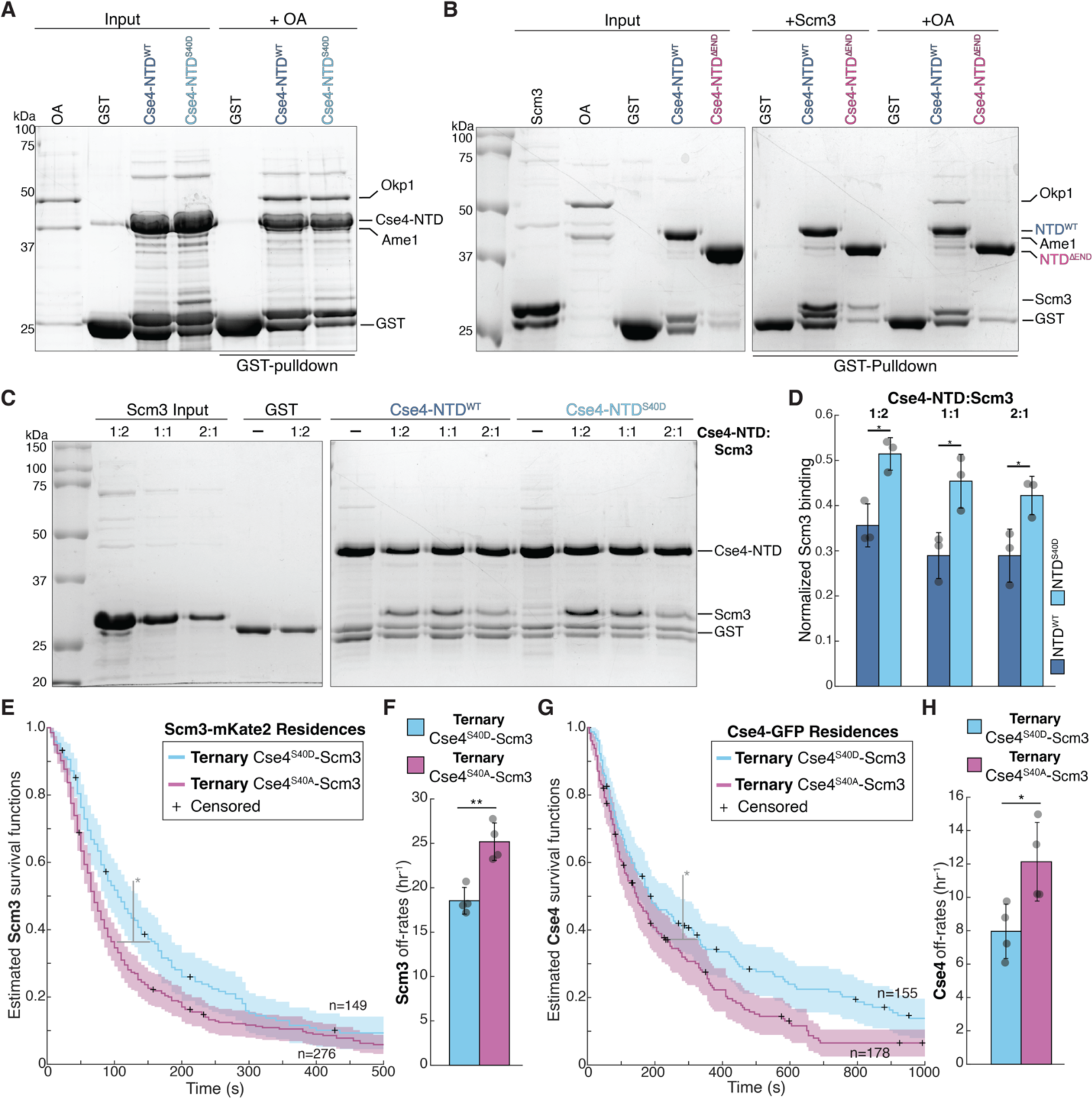
Cse4 END domain phosphorylation promotes Scm3 binding that stabilizes Cse4 on centromeric DNA. A. SDS-PAGE of GST pulldown assays of immobilized Cse4-NTD^WT^ and Cse4-NTD^S40D^ to test binding of OA. B. SDS-PAGE of GST pulldown assays of immobilized Cse4-NTD^WT^ and Cse4-NTD^ΔEND^ protein fusions binding to recombinant Scm3 and OA. C. SDS-PAGE of GST pulldown assays of immobilized Cse4-NTD^WT^ and Cse4-NTD^S40D^ to test binding of recombinant Scm3 at varying Scm3 concentrations. D. Quantification of Scm3 binding in pulldown assays normalized to Cse4 levels for NTD^WT^ (0.36 ± 0.05, 0.29 ± 0.05, 0.29 ± 0.06) and the NTD^S40D^ mutant (0.51 ± 0.04, 0.45 ± 0.06, 0.42 ± 0.04) at varying input concentrations (1:2, 1:1, and 1:2 Cse4-NTD: Scm3 respectively, experiment was repeated three times to generate averages, * indicates two-tailed *P*-value of .01 for 1:2, .02 for 1:1 and .03 for 2). E. Scm3 resides longer after colocalization with Cse4^S40D^ than with Cse4^S40A^. Estimated survival function plots of Kaplan–Meier analysis of the lifetimes in *CSE4-GFP SCM3-mKate2* (SBY22256) extract of Ternary^S40D^ Scm3 residences on CEN DNA (blue—median lifetime of 104 s, n = 149 over four experiments of ∼ 1,000 DNA molecules using different extracts) and Ternary^S40A^ Scm3 residences on CEN DNA (purple—median lifetime of 67 s, n = 276 over four experiments of ∼ 1,000 DNA molecules using different extracts). 95% confidence intervals indicated (shaded) right-censored lifetimes (plus icons) were included and with equivalent weighting in survival function estimates, * indicates significant difference two-tailed *P*-value of .01 as determined by log-rank test. F. Graph indicates quantification of the estimated off-rates of Ternary^S40D^ and Ternary^S40A^ Scm3 residences on CEN DNA (18.5 h^−1^ ± 2.3 h^−1^ and 25.2 ± 2.3 h^−1^ respectively, avg ± s.d. n = 395 over four experiments of ∼ 1,000 DNA molecules, ** indicates significant difference with two-tailed *P*-value of .004). G. Cse4^S40D^ resides longer after colocalization with Scm3 than Cse4^S40A^. Estimated survival function plots of Kaplan–Meier analysis of the lifetimes of Ternary^Scm3^ Cse4^S40D^ residences on CEN DNA (blue—median lifetime of 184 s, n = 155 over four experiments of ∼ 1,000 DNA molecules using different extracts) and Ternary^Scm3^ Cse4^S40A^ residences on CEN DNA (purple—median lifetime of 144 s, n = 178 over four experiments of ∼ 1,000 DNA molecules using different extracts). 95% confidence intervals indicated (shaded), right-censored lifetimes (plus icons) were included with equivalent weighting in survival function estimates, * indicates significant difference two-tailed *P*-value of .006 as determined by log-rank test. H. Graph indicates quantification of the estimated off-rates of Ternary^Scm3^ Cse4^S40D^ and Ternary^Scm3^ Cse4^S40A^ residences on CEN DNA (8.0 h^−1^ ± 1.6 h^−1^ and 12.1 ± 2.4 h^−1^ respectively, avg ± s.d. n = 322 over four experiments of ∼ 1,000 DNA molecules, * indicates significant difference with two-tailed *P*-value of .03).

It was previously reported that OA binding was enhanced when Scm3 was prebound to Cse4-NTD [27], so we tested whether prebound Scm3 could further stimulate OA binding in pulldown binding assays with Cse4-NTD^S40D^. Surprisingly, we did not detect a significant change in OA binding with either pre-Scm3 bound Cse4-NTD^WT^ or Cse4-NTD^S40D^ (**Supplemental Figure 3A**), possibly due to differences in assay conditions or protein constructs used for reconstitutions. To address this apparent discrepancy, we tested if OA recruitment was affected by the Cse4-S40D mutant in the context of the entire native Cse4 nucleosome instead of the just the N-terminus. To do this, we performed TIRFM endpoint colocalization assays to measure OA recruitment (using Ame1-GFP) in extracts containing either Cse4 or the phosphomimetic Cse4^S40D^ mutant. Consistent with the pulldown assays, Ame1-GFP colocalization levels were similar in extracts containing the Cse4^S40D^ mutant when compared to WT Cse4 (**Supplemental Figure 3B**). Together, these data indicate that the Cse4-S40D mutant specifically alters its interaction with Scm3 but not OA, providing a mutant to specifically explore the role of Scm3 binding to the Cse4 N-terminus.

### Scm3 binding to the Cse4 END promotes stable Cse4 recruitment

We asked if Scm3 binding to the Cse4 END stabilizes Cse4 on centromeres. First, we monitored the localization of phosphomimetic Cse4^S40D^ and phosphonull Cse4^S40A^ mutants in bulk kinetochore assembly assays. While Cse4^S40D^ assembled normally (**Supplemental Figure 4A** - middle), there was a reduction in Cse4^S40A^ centromeric association (**Supplemental Figure 4A** - right). To further characterize this defect, we performed TIRFM endpoint colocalization assays. Consistent with the bulk assembly assays, Cse4^S40D^-GFP localized to centromeric DNA similarly to WT (50% vs 55% respectively, **Supplemental Figure 4B**), while Cse4^S40A^-GFP recruitment was significantly reduced (28% vs 55% respectively, **Supplemental Figure 4B**). To further examine how the N-terminal interaction between Scm3 and Cse4 contributes to their stability on centromeric DNA, we performed time-lapse TIRFM colocalization assays. We simultaneously monitored orthogonally labeled Cse4^S40D^-GFP or Cse4^S40A^-GFP with Scm3-mKate2 on individual centromeric DNAs. We first asked how Scm3 stability is affected by Cse4 END domain phosphorylation within the context of the native nucleosome. Kaplan-Meier analysis of <u>Scm3 residences</u> on centromeric DNA that formed ternary colocalizations with either Cse4^S40D^ (Ternary Cse4^S40D^-Scm3) or Cse4^S40A^ (Ternary Cse4^S40A^-Scm3) revealed a significant increase in the median lifetime of Ternary Cse4^S40D^ when compared to Ternary Cse4^S40A^ Scm3 residences (104 s vs. 64 s, **Figure 5E**). Consistent with longer Ternary Cse4^S40D^-Scm3 lifetimes, off-rate analysis also revealed enhanced stability of Ternary Cse4^S40D^-Scm3 when compared to Ternary Cse4^S40A^-Scm3 (18.5 h^−1^ vs 25.2 h^−1^, **Figure 5F**). We next asked whether the prolonged Scm3 binding after ternary complex formation with Cse4^S40D^ also stabilized Cse4 by monitoring <u>Cse4 residences</u> on centromeric DNA. Kaplan-Meier analysis revealed that the median lifetimes of Ternary Cse4^S40D^-Scm3 residences were significantly increased compared to Ternary Cse4^S40A^-Scm3 residences on centromeric DNA (178 s vs. 144 s, **Figure 5G**). This was confirmed in off-rate analysis, which showed a significantly slower average off-rate of Ternary Cse4^S40D^-Scm3 versus Ternary Cse4^S40A^-Scm3 residences (18.5 h^−1^ vs 25.2 h^−1^, **Figure 5H**). Taken together, these analyses reveal that Cse4-S40 phosphorylation of the END region stabilizes both Scm3 and Cse4 association with centromeric DNA.

### Scm3 END binding can rescue defects in OA-dependent stabilization of Cse4

We next asked whether the contributions of Scm3 and OA to Cse4 centromere retention are independent. First, we tested whether the Cse4-L41 mutants that alter OA:END association affect Scm3 binding. We found that neither the Cse4-L41A or Cse4-L41D mutant affected Scm3 association to the Cse4-NTD in pulldown assays (**Supplemental Figure 5)**, confirming that *cse4-L41* mutants specifically affect OA binding. We next asked whether the *cse4^L41A^* mutant could be suppressed by enhanced Scm3 binding via the Cse4^S40D^ phosphomimetic mutation. We generated a *cse4^S40D,L41A^* double mutant and performed bulk assembly assays. Cse4 levels were restored on centromeric DNA in the *cse4^S40D,L41A^* mutant compared to *cse4^L41A^* (**Supplemental Figure 6A**). OA association was not restored, confirming that enhanced Scm3 binding to the END can suppress defects in Cse4 recruitment due to a lack of END:OA interaction. We quantified the rescue in TIRFM endpoint colocalization assays and found that Cse4^S40D,L41A^*-*GFP levels were restored relative to the Cse4^L41A^*-*GFP mutant, while the levels of Ame1-mKate2 remained disrupted in both mutants (**Figure 6A, B**). In a complementary experiment, we asked whether the Cse4-S40D mutant could rescue the Cse4 centromeric localization defect in the *ame1^I195Y^* mutant that disrupts OA interaction with the Cse4 END. The Cse4^S40D^-GFP localization was significantly rescued when compared to WT Cse4-GFP in Ame1^I195Y^ mutant extracts (**Figure 6C, D**).

**Figure 6.**
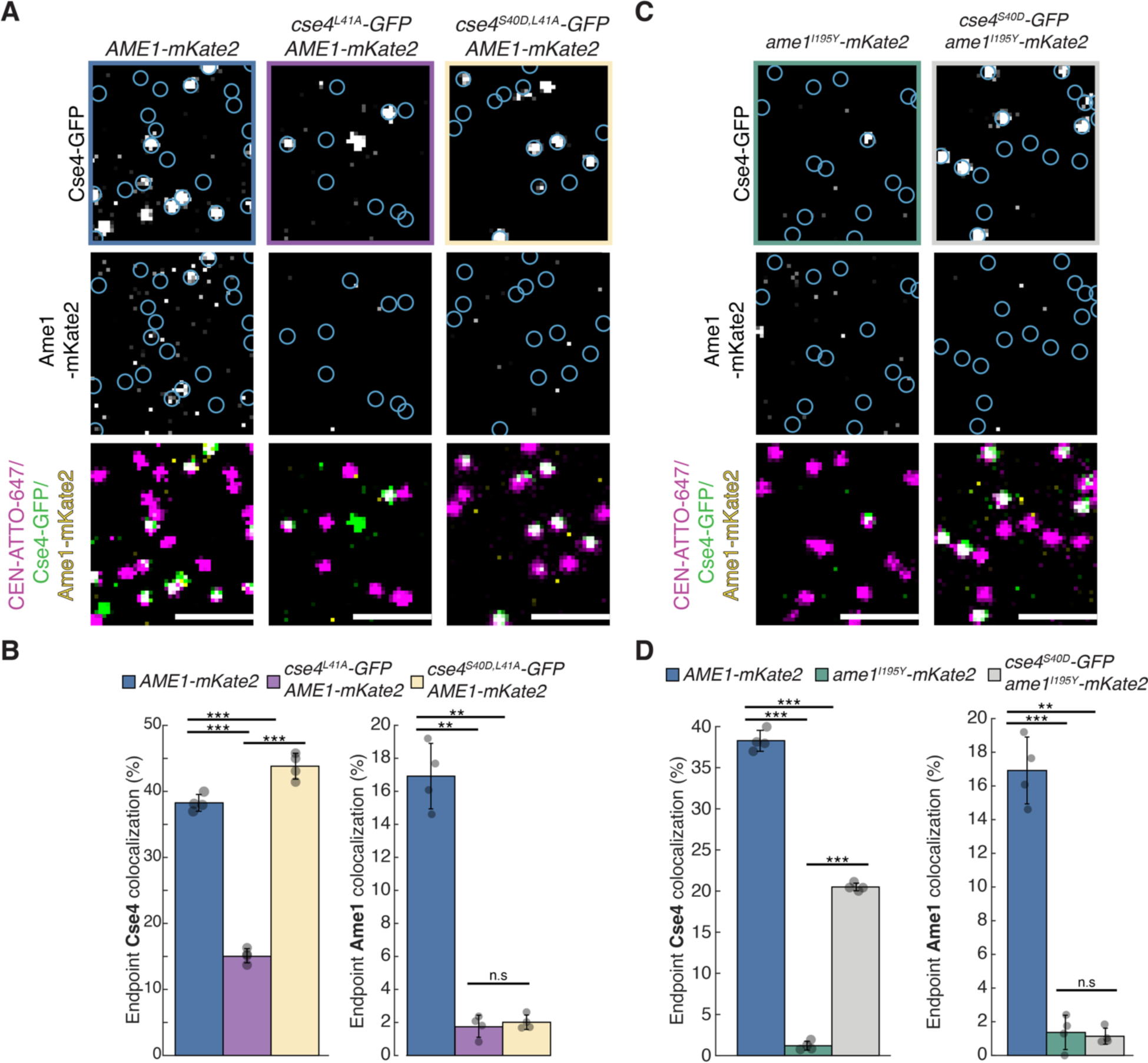
Enhanced Scm3 binding to the Cse4 END rescues defects caused by OA disruption. A. Example images of TIRFM endpoint colocalization assays. Top panels show visualized Cse4-GFP on CEN DNA in *CSE4-GFP Ame1-mKate2* (SBY22244) extracts (top-left panel), *cse4^L41A^-GFP AME1-mKate2* (SBY22931) extracts (top-middle panel), or *cse4^S40D/L41A^-GFP AME1-mKate2* (SBY22929) extracts (top-right panel) with colocalization shown in relation to identified CEN DNA in blue circles. Bottom panels show overlay of CEN DNA channel (magenta) with Cse4-GFP (green) and Ame1-mKate2 (yellow), scale bars 2 μm. B. Graph indicates quantification of endpoint colocalization of Cse4-GFP with CEN DNA in extracts from *CSE4-GFP AME1-mate2*, *cse4^L41A^-GFP AME1-mKate2* or *cse4^S40D,L41A^-GFP AME1-mKate2* genetic backgrounds (38 ± 1.3%, 15 ± 2.3%, 44 ± 2.0%, avg ± s.d. n = 4 experiments, each examining ∼ 1,000 DNA molecules from different extracts, *** indicates significant difference between *CSE4-GFP AME1-mKate2* and *cse4^L41A^-GFP AME1-mKate2* with two-tailed *P*-value of 1.6E-5 and between *cse4^S40D,L41A^-GFP AME1-mKate2* and *cse4^L41A^-GFP AME1-mKate2* with two-tailed *P*-value of 1.3E-6, ** indicates significant difference between *CSE4-GFP AME1-mKate2* and *cse4^S40DL,41A^-GFP AME1-mKate2* with two-tailed *P*-value of 0.005.), or colocalization of Ame1-mKate2 with Cse4-GFP (17 ± 2.0%, 1.7 ± 0.4%, 2 ± 0.4%, avg ± s.d. n = 4 experiments, each examining ∼ 1,000 DNA molecules from different extracts, ** indicates significant difference between *CSE4-GFP AME1-mKate2* and *cse4^L41A^-GFP AME1-mKate2* with two-tailed *P*-value of 6.4E-4 and *CSE4-GFP AME1-mKate2* and *cse4^S40D,L41A^-GFP AME1-mKate2* with two-tailed *P*-value of 6.8E-4, n.s. indicates no significant difference between *cse4^S40D,L41A^-GFP AME1-mKate2* and *cse4^L41A^-GFP AME1-mKate2* with two-tailed *P*-value of 0.3). C. Example images of TIRFM endpoint colocalization assays. Top panels show visualized Cse4-GFP on CEN DNA in *CSE4-GFP ame1^I195Y^* (SBY23105 - top-left panel), or *cse4^S40D^-GFP ame1^I195Y^* extracts (SBY23163 - top-right panel) with colocalization shown in relation to identified CEN DNA in blue circles. Bottom panels show overlay of CEN DNA channel (magenta) with Cse4-GFP (green), scale bars 2 μm. D. Graph indicates quantification of Cse4-GFP endpoint colocalization with CEN DNA in *CSE4-GFP*, *CSE4-GFP ame1^I195Y^* or *cse4^S40D^-GFP ame1^I195Y^* extracts (38 ± 1.3%, 1 ± 0.5%, 21 ± 0.5%, avg ± s.d. n = 4 experiments, each examining ∼ 1,000 DNA molecules from different extracts, *** indicates significant difference between *CSE4-GFP AME1-mKate2* and *CSE4-GFP ame1^I195Y^-mKate2* with two-tailed *P*-value of 7.0E-7, between *CSE4-GFP AME1-mKate2* and *cse4^S40D^-GFP ame1^I195Y^-mKate2* with two-tailed *P*-value of 1.2E-5 and between *cse4^S40D^-GFP ame1^I195Y^-mKate2* and *CSE4-GFP ame1^I195Y^-mKate2* with two-tailed *P*-value of 2.7E-9.) or colocalization of Ame1-mKate2 with Cse4-GFP (17 ± 2.0%, 1.4 ± 1.0%, 1.1 ± 0.5%, avg ± s.d. n = 4 experiments, each examining ∼ 1,000 DNA molecules from different extracts, *** indicates significant difference between *CSE4-GFP AME1-mKate2* and *CSE4-GFP ame1^I195Y^-mKate2* with two-tailed *P*-value of 1.3E-4 and *CSE4-GFP AME1-mKate2* and *cse4^S40D^-GFP ame1^I195Y^-mKate2* with two-tailed *P*-value of 5.9E-4, n.s. indicates no significant difference between *cse4^S40D^-GFP ame1^I195Y^-mKate2* and *CSE4-GFP ame1^I195Y^-mKate2* with two-tailed *P*-value of 0.7).

To ensure that the rescue of Cse4 localization in the Cse4-S40D mutant did not rely on transient OA binding or other OA activity, we asked if Cse4^S40D^ could still be stabilized on centromeres in the complete absence of OA. We utilized a previously described auxin-inducible degron system to rapidly degrade Okp1 (*okp1-AID)* in cells containing Cse4-GFP or Cse4^S40D^-GFP prior to generating extracts. We confirmed that OA depletion severely disrupted Cse4-GFP localization and it was rescued by the Cse4^S40D^-GFP mutant in endpoint assays (**Supplement Figure 6B**). However, Cse4 levels were not rescued to the same extent as in the Cse4^S40D,L41A^ mutant (**Figure 6B, C**), likely because Cse4^L41A^ retains residual OA binding (**Figure 3A**). Taken together, our results suggest that phosphorylation of Cse4 END at S40, which stimulates Scm3 binding, can promote retention of Cse4 at centromeres independently from OA binding to Cse4 END, but that OA and Scm3 binding are both needed for full centromere localization.

### OA and Scm3 contribute to Cse4 centromeric localization in vivo

We next analyzed the cellular consequences of the dual contributions of Scm3 and OA binding to the Cse4 END domain. Consistent with a previous report [32], we found that the *cse4^L41A^*mutant exhibited a significant growth defect at higher temperatures (**Figure 7A**). Strikingly, the growth defect was largely rescued in the *cse4^S40D,L41A^* mutant (**Figure 7A**). To better understand the cause of the *cse4^L41A^* mutant growth defect, we first assayed cell cycle progression at high temperature. *CSE4*, *cse4^L41A^* and *cse4^S40D,L41A^* mutant cells were released from a G1 arrest to 36 degrees and monitored for cell cycle progression via budding index. WT cells were large-budded 90’ after release and completed cell division within 150 min (**Figure 7B**). The *cse4^S40D,L41A^*mutant cells progressed more slowly and became large-budded 120’ after release but eventually completed cell division. In contrast, *cse4^L41A^* cells accumulated as large-budded cells and never divided, suggesting activation of the spindle checkpoint (**Figure 7B**). To test this, we crossed the *cse4^L41A^* mutant to a *mad1Δ* checkpoint deletion mutant and found the double mutants were inviable (**Figure 7C**). These data indicate that the spindle checkpoint is required for the viability of *cse4^L41A^* mutant cells, strongly suggesting that there are defective kinetochore-microtubule attachments.

**Figure 7.**
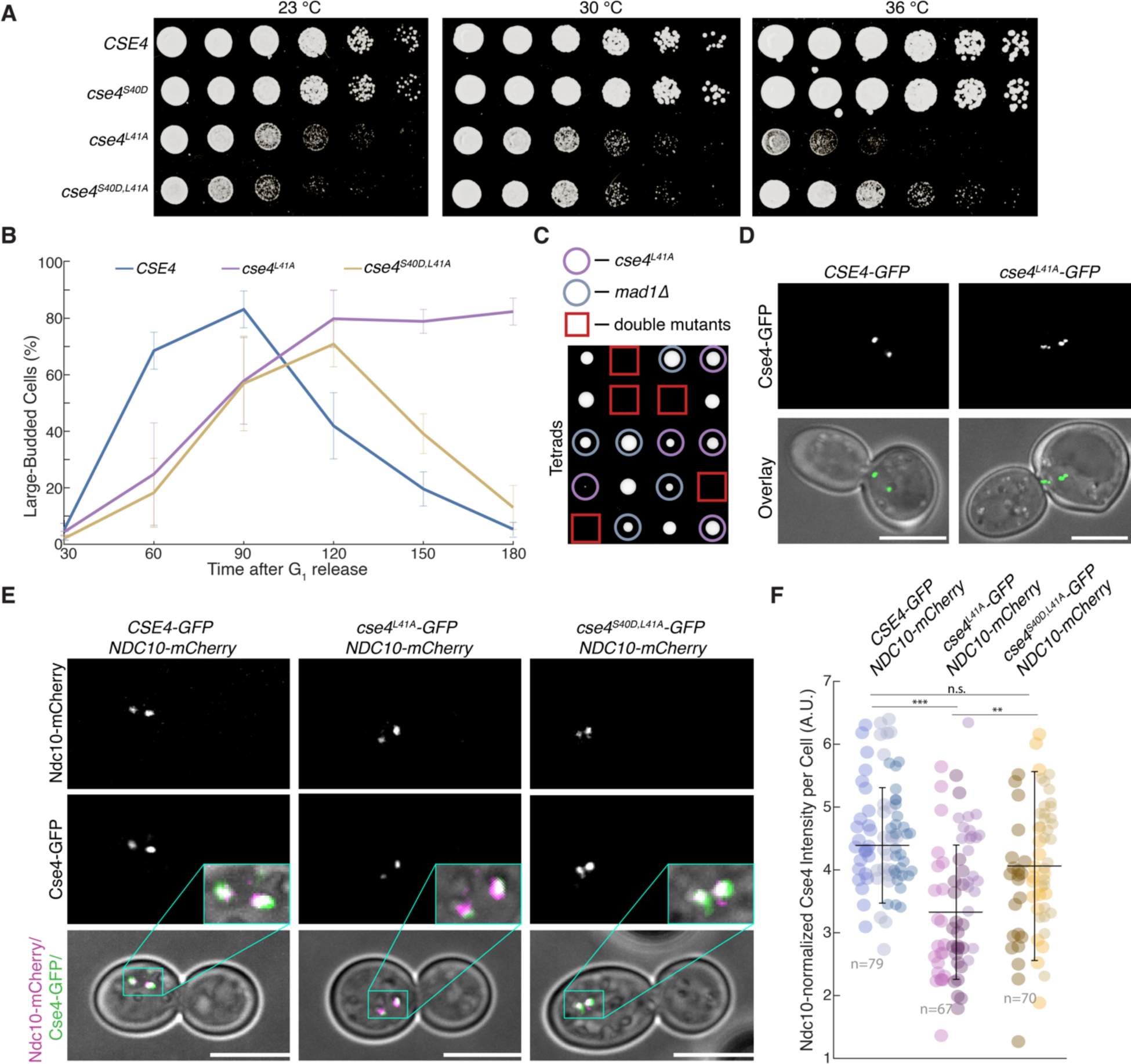
Cse4-S40D can suppress reduced Cse4 centromeric localization caused by disruption of OA binding to END. A. Five-fold serial dilutions of *CSE4-GFP* (*SBY21863*), *cse4^S40D^-GFP* (*SBY20017*), *cse4^L41A^-GFP* (*SBY22811*) and *cse4^S40D,L41A^-GFP* (*SBY22914*) were plated and grown on YPD for two days at 23°, 30° and 36°. B. *Cse4-L41A* mutant causes mitotic arrest at high temperatures. *CSE4-GFP* (black), *cse4^L41A^-GFP* (green) and *cse4^S40D,L41A^-GFP* (magenta) cells were released from G_1_ and fixed every 30 min, and the proportion of large-budded cells quantified. Each time point represents the percentage of budded cells for each strain (about 200 cells, avg. ± s.d., n = 3 biological replicates). C. Haploid progeny from sporulation of *cse4^L41A^* (*SBY22811*)/*mad1τ<* (*SBY22811*) heterozygous diploid, red square indicates double mutant haploid spores. Tetrads are aligned horizontally. D. Representative images of *CSE4-GFP* (left) and *cse4^L41A^-GFP* (right) cells 90 min after G_1_ release. Scale bars 5 μm. E. Example fluorescence microscopy images of *CSE4-GFP NDC10-mCherry* (SBY21973 - left), *cse4^L41A^-GFP NDC10-mCherry* (SBY23101 - middle) and *cse4^S40D,L41A^-GFP NDC10-mCherry* (SBY23474 - right) cells showing visualized Ndc10-mCherry (top panels), Cse4-GFP (middle panels) and overlay of Ndc10-mCherry (magenta) and Cse4-GFP (green) on plane polarized illumination of cell. Expanded region around kinetochores highlighted (middle panel inset). Scale bars 5 μm. F. Graph indicates quantification of normalized centromere-associated Cse4-GFP intensity per cell of *CSE4-GFP NDC10-mCherry* (left) *cse4^L41A^-GFP NDC10-mCherry* (middle) and *cse4^S40D,L41A^-GFP NDC10-mCherry* (right) cells (4.4 ± 0.9 a.u., 3.3 ± 1.1 a.u., 4.1 ± 1.5 a.u., avg ± s.d. n = 3 experiments, each examining ∼ 25 cells). * Indicates significant difference and n.s. indicates non-significant as determined by t-Test (WT:L41A *P*-value of 1.6E-9, WT:DA *P*-value of 0.12 and L41A:DA *P*-value of 0.001). Each spot represents calculated intensity for one cell, different colors indicate biological replicates.

To further examine the potential kinetochore defects in the *cse4^L41A^*cells, we imaged Cse4-GFP in the cells. In yeast, kinetochores cluster into two foci due to the pulling forces of microtubules on bioriented kinetochores [38–40]. As expected, wild type Cse4-GFP exhibited two foci in the large-budded cells at metaphase (**Figure 7D** - left). In contrast, there were multiple Cse4^L41A^-GFP foci, suggesting aberrant kinetochore attachments (**Figure 7D** - right). To determine if the defective kinetochore localization was related to Cse4 stability at centromeres, we monitored Cse4 levels at centromeres relative to the fiducial centromere marker Ndc10-mCherry after shifting cells to high temperature (36 °C) under mitotic arrest for 2 hrs (**Figure 7E**). Kinetochore clusters were identified via Ndc10 and used to quantify Cse4 centromeric intensity relative to Ndc10 on a per cluster basis and then summed per cell for comparison. Strikingly, Cse4 levels at centromeres were significantly reduced in the *cse4^L41A^* mutant and rescued to normal levels in the *cse4^S40D,L41A^* double mutant (**Figure 7E, F**). We used the same method to analyze Cse4 levels in cells depleted of OA (*okp1-*AID) and confirmed that Cse4 levels at centromeres were significantly reduced (**Supplemental Figure 7A, B**). We also monitored OA levels using GFP-tagged Ame1 under the same conditions in *cse4^L41A^* and *cse4^S40D,L41A^* cells. Consistent with the *in vitro* assays, the centromere-associated levels of Ame1-GFP were significantly reduced in the *cse4^L41A^*mutant and were not rescued in *cse4^S40,L41A^* mutant cells (**Supplemental Figure 8**). Together, these data show that END domain contributes to Cse4 centromere localization *in vivo*.

## DISCUSSION

### Stable Cse4 centromeric localization requires its END domain

While recent structural models have provided key insights into overall inner kinetochore architecture, the role of the disordered Cse4-NTD in kinetochore assembly remained unclear because it is not visible within the structures of the current nucleosome reconstitutions. Using single molecule fluorescence microscopy, we discovered that the Cse4 END region is critical for Cse4 centromeric localization. To determine whether the nucleosome assembly process in conventional biochemical reconstitutions differed from *de novo* assembly with native components, we tested the modified centromere sequence (C0N3) used in reconstitutions in our TIRFM assay. We observed a marked reduction in Cse4 centromeric localization by the C0N3 template that replaces CDEII with a strong nucleosome positioning sequence when compared to the native centromere sequence of CEN DNA templates. These data suggest there are additional assembly pathways or contributing interactions that play a significant role in formation of the centromeric nucleosome. These factors may be more easily distinguished in reconstitutions using cell extracts that contain these additional and potentially previously unidentified kinetochore assembly components. Our *de novo* kinetochore assembly assays are therefore a powerful tool to dissect pathways that are not easily detected by other assays. Using this approach, we discovered that the Cse4 END region is critical for Scm3 binding in addition to its previously established essential binding interaction with OA, and that both END interactions have a critical role in stabilizing the centromeric nucleosome.

### OA and Scm3 stabilize Cse4 via the END

Cse4 END binding to OA has been well characterized and its essential function was assumed to be OA recruitment to the kinetochore. However, we found that even modest disruption of OA binding to the Cse4 END (via Cse4^L41A^) significantly disrupted both Cse4 and OA centromeric localization in vitro. Consistent with this, OA also contributes to Cse4 centromeric levels *in vivo*. The *cse4^L41A^* mutant that weakens OA binding arrests in mitosis at high temperature due to spindle checkpoint activation and exhibits reduced Cse4 centromere levels. In addition, analysis of centromeric chromatin structure in vivo showed that the nucleosome structure was disrupted in the absence of OA. Together, these results suggest that OA binding to the Cse4 END domain not only recruits OA to the kinetochore, but also stabilizes the centromeric histone in the process. This END-dependent role of maintaining centromeric localization of Cse4 is consistent with the role of the N-terminal tail of CENP-A in *C. elegans,* where it was shown to be required for proper centromeric assembly [34, 35]. In the future, it will be important to determine whether OA’s interactions with additional kinetochore proteins in the CCAN further contribute to Cse4 stability at the centromere.

We also identified a role for the Scm3 chaperone in Cse4 localization via the END domain. We previously discovered that Scm3 stabilized Cse4 on centromeric DNA, and here we show it is at least partially through Scm3 binding to the Cse4 END, similar to the role of recruitment of KNL-2 by the N-terminal tail of CENP-A in *C. elegans* [34, 35]. Ipl1 phosphorylation of Cse4-S40 promotes Scm3 binding and this END-dependent regulation of Scm3 binding is independent of OA. The distinct roles of Scm3 and OA in stabilizing the Cse4 nucleosome via the same END region suggests these activities might be temporally separated. Scm3 has much higher affinity for the Cse4 histone fold domain than the N-terminus, so we presume it initially targets Cse4 to centromeres via this interaction. DNA wrapping would displace Scm3 from the Cse4 histone fold domain, making it accessible to bind to the N-terminus after nucleosome formation. The dual Scm3 binding sites on Cse4 may explain its exchange at kinetochores throughout the cell cycle [4]. Although OA binds to the Cse4 END with higher affinity than Scm3, it is unclear whether they simultaneously bind. The temporal order of Scm3 and OA roles in stabilizing Cse4 during kinetochore assembly is open to further study.

### Scm3 binding to the Cse4 END can compensate for weakened OA recruitment

The enhanced Scm3 binding to the Cse4 END due to Cse4-S40 phosphorylation rescued defects in Cse4 centromeric localization caused by the Cse4^L41A^ OA binding mutant both in vitro and in vitro. This rescue was specifically driven by Scm3 binding as OA levels were not rescued in the Cse4^S40D,L41A^ double mutant. Moreover, the Cse4^S40D^-dependent rescue in Cse4 localization could be maintained even in the most extreme case, where OA was absent during kinetochore assembly (**Supplemental Figure 6B**). Together, these results highlight previously unknown functions for Scm3 in promoting Cse4 stability on centromeric DNA, furthering our understanding of the unique Scm3 chaperone protein which is part of a larger family of histone chaperones whose set of cellular functions continue to expand [41]. Moreover, these additional Scm3 functions depend on interactions at the kinetochore that are not via the histone fold domain, suggesting additional chaperone functions important for kinetochore assembly.

### Regulation of Cse4 localization

Our finding that Ipl1 phosphorylation of Cse4-S40 promotes Cse4 localization highlights a previously unknown regulatory point of kinetochore assembly. While it had been previously shown that Ipl1 phosphorylation of Cse4 played a role in chromosome segregation [29], the underlying mechanisms were not known. Here we identified the function of one of the four previously characterized sites (S22, S33, S40 and S105). We note that this phosphorylation event is not essential, likely due to the multiple mechanisms we’ve identified that ensure Cse4 localization and stability *in vivo*. We propose that S40 phosphorylation occurs directly at centromeres because Ipl1 associates with the Ndc10 protein that binds to the centromere. Although Ipl1 is most well-known for its role in error correction, our work has identified another function for Ipl1 in kinetochore assembly. It was previously established that Ipl1 facilitates kinetochore assembly by relieving autoinhibition of the interaction between Mif2 and the MIND complex by phosphorylation the Dsn1 protein [42, 43]. The Ipl1 regulation of Dsn1 is a conserved kinetochore assembly mechanism, and it will be interesting to determine whether the regulation of the centromeric histone variant is conserved. CENP-A is known to be phosphorylated by Aurora B in other organisms [44], but it isn’t yet clear whether the chaperone HJURP binds to a second region in CENP-A.

It is perplexing that the centromere is a poor nucleosome assembly template in conventional biochemical reconstitutions. The inherent instability of the yeast centromeric nucleosome has precluded kinetochore reconstitutions using complete native centromere sequences. Here, we have identified multiple previously unknown mechanisms that ensure the stabilization of the yeast centromeric nucleosome. We propose that yeast use a multi-step “licensing” assembly mechanism to ensure that chromosomes assemble a single centromeric nucleosome (**Figure 8**). We presume that the initial Cse4 association with centromeres is driven by Scm3 binding to the CATD within the histone fold domain to target Cse4 to the centromere. Once Cse4 is at the centromere, Ipl1 would be in the proximity of the Cse4 END and phosphorylate it to promote Scm3 binding to the N-terminus. The END domain would also bind to OA, which would further stabilize the centromeric nucleosome. Because Ipl1 is associated with the CBF3 complex that exclusively binds to the CDEIII centromere element, this licensing could not occur in euchromatin and would therefore help prevent stable ectopic Cse4 deposition to maintain chromosomal integrity during mitosis. This complex regulatory schema relies on the unique Cse4 END region, which may be a consequence of the specific point centromere architecture present in budding yeast where a single centromeric nucleosome is sufficient to assemble the kinetochore. The stringent centromere sequence composition requires coordination of several kinetochore proteins to form centromeric nucleosomes, ensuring the specificity of centromeric histone deposition which is crucial to cell division and organismal survival. Further study of kinetochore assembly pathways under differing cellular contexts is critical for understanding how kinetochores form and function in cells.

**Figure 8.**
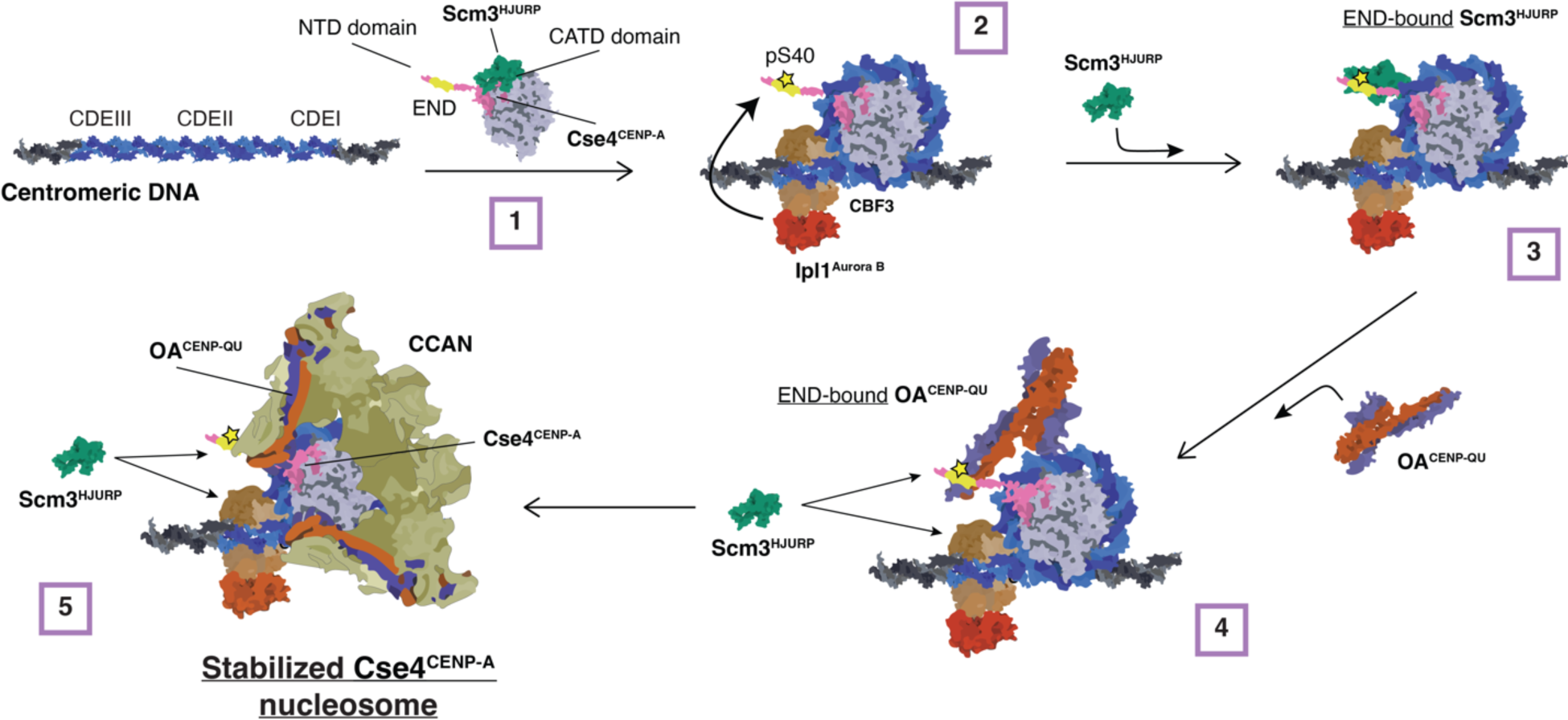
Model for stabilization of Cse4 at centromeres. Schematic diagram of proposed model of Cse4 nucleosome formation that relies on END recruitment of the essential chaperone Scm3 and the essential CCAN component OA for stability prior to kinetochore assembly. 1. CATD-bound Scm3 targets Cse4 to the centromere. 2. CATD-bound Scm3 is released when DNA wraps to form nucleosome and centromere-associated (CBF3) Ipl1 phosphorylates Cse4 at S40. 3. Scm3 binds to NTD, promoted by END phosphorylation at S40, to initially stabilize Cse4 nucleosome. 4. OA is recruited to END to further stabilize Cse4 and recruit remaining CCAN components while Scm3 remains in exchange at centromere. 5. Upon full recruitment of CCAN, Cse4 nucleosome is fully stabilized while Scm3 remains in exchange at centromere.

## METHODS

### Yeast Methods

The *S. cerevisiae* strains used in this study are listed in Supplemental Table 1 and are derived from SBY3 (W303). Strains were generated using standard genetic crosses, media and PCR-based tagging techniques [45–47]. The plasmids and primers used to generate strains are listed in Supplemental Tables 2 and 3, respectively. Single point mutants (*cse4-S40D*, *cse4-S40A*, and *cse4-L41A*) were introduced into endogenous alleles using standard yeast CRISPR techniques [48]. *CSE4-GFP* and all mutant derivatives were internally tagged at residue 80 with eGFP including linkers on either side (pSB1617) and then expressed from its native promoter at the endogenous locus. *CSE4-mCherry* is equivalently constructed with mCherry in place of eGFP (pSB3271) and expressed similarly (SBY20858). Endogenous genes that were altered to include fluorescent protein alleles (eGFP, mCherry or mKate2) tags or auxin-inducible degrons (-IAA7) were constructed by standard PCR-based integration techniques [45] and confirmed by PCR. All liquid cultures were grown in yeast peptone dextrose rich (YPD) media. For kinetochore assembly assays, cells were arrested in mitosis by diluting log phase cultures into benomyl-containing liquid media to a final concentration of 30 μg/mL and grown for another three hours until at least 90% of cells were large-budded. For strains with auxin inducible degron (AID) alleles (*ipl1-AID*, *okp1-AID*), cultures were treated with 500 μM indole-3-acetic acid (IAA, dissolved in DMSO) for the final 60 min of mitotic arrest as described previously [49–51]. For strains with galactose inducible alleles (*pGAL-SCM3*), cultures were grown in raffinose and mitotically arrested in raffinose and benomyl-containing liquid media and then treated with 4% galactose for the final 60 min of mitotic arrest. Growth assays were performed by diluting log phase cultures to OD600 ∼ 1.0 from which a 1:5 serial dilution series was made and plated on YPD and incubated at 23 °C, 30 °C and 36 °C. For budding index assays, log phase liquid cultures were arrested with alpha-factor for three hours at room temperature and then released and temperature shifted to 36 °C. Cells were collected and fixed every 20 mins and alpha factor was added once ∼50% cells appeared large budded to prevent further division. Cells were visualized on concanavalin A-functionalized coverslips using plane polarized light with a 60X objective on an inverted Nikon light microscope. At each respective time point and for each respective strain, ∼200 cells were counted for large-budded appearance.

### Whole Cell Extract preparation for kinetochore assembly assays

For kinetochore assembly assays, cells were grown in liquid YPD media to log phase and arrested in mitosis in 500 mL total volume and then harvested by centrifugation. All subsequent steps were performed on ice with 4 °C buffers. Cells were washed once with dH2O with 0.2 mM PMSF, then once with Buffer L (25 mM HEPES pH 7.6, 2 mM MgCl2, 0.1 mM EDTA pH 7.6, 0.5 mM EGTA pH 7.6, 0.1 % NP-40, 175 mM K-Glutamate, and 15% Glycerol) supplemented with protease inhibitors (10 mg/ml leupeptin, 10mg/ml pepstatin, 10mg/ml chymostatin, 0.2 mM PMSF) and 2 mM DTT. Cell pellets were then snap frozen in liquid nitrogen and lysed using a Freezer/Mill (SPEX SamplePrep), using 10 rounds that consisted of 2 min of bombarding the pellet at 10 cycles per second, then cooling for 2 min. The subsequent powder was weighed and then resuspended in Buffer L according to the following calculation: weight of pellet (g) x 1.5 =number of mL of Buffer L. Resuspended cell lysate was thawed on ice and clarified by centrifugation at 16,100 g for 30 min at 4 °C, the protein-containing layer was extracted with a syringe and then aliquoted and snap frozen in liquid nitrogen. The resulting soluble whole cell extracts (WCE) generally had a concentration of 60–80 mg/mL. The pellets, powder, and WCE were stored at −80 °C.

### Preparation of DNA templates and Dynabeads for assembly assays

As described previously [22], plasmid pSB963 was used to generate the WT CEN DNA templates and pSB972 was used to generate the CEN^mut^ template used in this study. PCR products were purified using the Qiagen PCR Purification Kit. In the case of bulk assembly assays, purified CEN DNA was conjugated to Streptadivin-coated Dynabeads (M-280 Streptavidin, Invitrogen) for 2.5 hr at room temperature, using 1 M NaCl, 5 mM Tris-HCl (pH7.5), and 0.5 mM EDTA as the binding and washing buffer. For single molecule TIRFM assays, purified CEN DNA was diluted in TE to a final concentration ∼100 pM.

### Bulk assembly assays and immunoblotting

Bulk kinetochore assembly was performed with whole cell extract and WT CEN or CEN^mut^ DNA conjugated to Streptadivin-coated Dynabeads as previously described [49]. Briefly, 0.5 mL of whole cell extract and 25 μl of DNA coated M280 Dynabeads (250bp WT CEN) were incubated at room temperature for either 90 min to allow complete kinetochore assembly or incubated at room temperature and stopped at 20 min, 40 min and 80 min by washing 3x and then resuspension in 1x Laemmli sample buffer (Bio-Rad). Samples were then resolved via 10% SDS-PAGE. Proteins were then transferred from SDS-PAGE gels onto 0.22 μM cellulose paper, blocked at room temperature with 4% milk in PBST, and incubated overnight at 4 °C in primary antibody. Antibody origins and dilutions in PBST were as follows: α-Cse4 (9536 [52]; 1:500), α-Ndc10 (OD1 [14]; 1:5000, α-Ndc80 (OD4 [50]; 1;10000.The anti-Mif2, anti-Ctf19, anti-Okp1 anti-Ame1 antibodies were generated in rabbits against their respective recombinant protein by Genscript. The company provided affinity-purified antibodies for each protein that we validated by immunoprecipitation of the respective target protein from yeast strains with an endogenously V5-tagged (Mif2, Ame1) or 13myc (Ctf19, Okp1) allele of the target protein, followed by confirmation of antibody recognition a protein of the correct molecular weight that was also recognized by α-V5 antibody (Invitrogen; R96025; 1:5000) or α-myc antibody (Cell Signaling Technology; 71D10, 1:10000). We subsequently used the antibodies at a the following dilutions: Mif2 (3747; 1:3000), Ctf19 (498; 1:5000), Ame1 (2181; 1:5000), Okp1 (28; 1;5000). Secondary antibodies were validated by the same methods as the primary antibodies as well as with negative controls lacking primary antibodies to confirm specificity. Blots were then washed again with PBST and incubated with secondary antibody at room temperature. Secondary antibodies were α-mouse (NA931) or α-rabbit (NA934), horseradish peroxidase-conjugated purchased from GE Healthcare and used at 1:1000 dilution in 4% milk in PBST. Blots were then washed again with PBST and ECL substrate (Thermo Scientific) was used to visualize the proteins on a ChemiDoc Imager (Bio-Rad). Uncropped and unprocessed scans of blots are provided in the Source Data file.

### Recombinant protein expression and purification

All constructs were expressed in BL21 Rosetta2 cells (EMD Millipore) as follows: cells were grown in TB media to an O.D. of ∼0.6 and then induced with 0.1M IPTG at 18 °C for 16 hrs. After induction cells were harvested then re-suspended in binding buffer (25mM HEPES, 175 mM KkGlut, 6 mM Mg(OAc)_2_, 15% glycerol, and 0.1% NP40, pH 7.6) supplemented with protease inhibitors and then lysed via sonication. After which, extract debris was pelleted at 14k RCF for 45 min. For HIS-tagged constructs (Scm3, OA), clarified cell supernatant was supplemented with 10mM immidazole and then incubated with HisPur cobalt resin (Thermo Fisher Scientific) for 40 min at 4 °C and then washed with 20x resin bed volume of binding buffer supplemented with 20mM immidazole. Proteins were then eluted with binding buffer supplemented with 150mM immidazole and fractions were analyzed via SDS-PAGE. The purest elution fractions containing were pooled, and buffer-exchanged to remove immidazole. Protein aliquots were snap frozen in liquid nitrogen. For GST-tagged proteins (Cse4-NTD constructs), clarified cell supernatant was incubated with Glutathione MagBeads (GeneScript) for 40 min at 4 °C and then washed with 20x bead volume of binding buffer. Washed beads were then analyzed via were analyzed via SDS-PAGE and diluted in binding buffer to a final concentration of ∼150 μM. Protein aliquots were snap frozen in liquid nitrogen.

### In vitro binding assays

GST-Cse4 NTD constructs (WT, S40D, L41A, L41D, ΔEND) were diluted to ∼15 μM in binding buffer (25mM HEPES, 175 mM KkGlut, 6 mM Mg(OAc)_2_, 15% glycerol, and 0.1% NP40, pH 7.6) with ∼15 μM of Ame1-6xHIS/Okp1 or Scm3-6xHIS and then mixed via rotation at 25 °C for 30 min. Resin was then washed at least three times with binding buffer to remove unbound protein, and bound protein then eluted with Laemmli Sample Buffer and boiling. Input and bead-bound proteins were analyzed by SDS–PAGE visualized by Coomassie brilliant blue staining. For Scm3 binding titrations of GST-Cse4 NTD (WT and S40D), GST constructs were diluted to ∼15 μM in binding buffer containing 30 μM, 15 μM or 7.5 μM Scm3-6xHIS (2:1, 1:1, 1:2 Cse4-NTD:Scm3 respectively) and then mixed via rotation at 25 °C for 30 min. Resin was then washed and analyzed by SDS–PAGE as before. To determine relative binding of Scm3 to Cse4-NTD constructs, SDS PAGE Scm3-6xHIS protein band total intensity was determined using FIJI [53] and normalized to total intensity of Cse4-NTD protein band in corresponding lane. This experiment was repeated three times and binding ratios were averaged between repeats of input concentrations.

### Single molecule TIRFM slide preparation

Coverslips and microscope slides were ultrasonically cleaned and passivated with PEG as described previously [54, 55]. Briefly, ultrasonically cleaned slides were treated with vectabond (Vector Laboratories) prior to incubation with 1% (w/v%) biotinylated mPEG-SVA MW-5000K/mPEG-SVA MW-5000K (Lysan Bio) in flow chambers made with double-sided tape. Passivation was carried out overnight at 4 °C. After passivation, flow chambers were washed with Buffer L and then incubated with 0.3 M BSA/0.3M Kappa Casein in Buffer L for 5 min. Flow chambers were washed with Buffer L and then incubated with 0.3M Avidin DN (Vector Laboratories) for 5 min. Flow chambers were then washed with Buffer L and incubated with ∼100 pM CEN DNA template for 5 min and washed with Buffer L. For endpoint colocalization assays, slides were prepared as follows: Flow chambers were filled with 100 μL of WCE containing protein(s) of interest via pipetting and wicking with filter paper. After addition of WCE, slides were incubated for 90 min at 25°C and then WCE was washed away with Buffer L. Flow chambers were then filled with Buffer L with oxygen scavenger system [56] (10 nM PCD/2.5 mM PCA/1mM Trolox) for imaging. For time-lapse colocalization assays, slides were prepared as follows: On the microscope, 100 μL WCE spiked with oxygen scavenger system was added to flow chamber via pipetting followed by immediate image acquisition.

### Single molecule TIRFM colocalization assays image collection and analysis

All images were collected on a Nikon TE-2000 inverted RING-TIRF microscope with a 100x oil immersion objective (Nikon Instruments) with an Andor iXon X3 DU-897 EMCCD camera. Images were acquired at 512 px x 512 px with a pixel size of 0.11 µm/px at 10MHz. Atto-647 labeled CEN DNAs were excited at 640 nm for 300 ms, GFP-tagged proteins were excited at 488 nm for 200 ms, and mCherry/Mkate2-tagged proteins were excited at 561 nm for 200 ms. For endpoint colocalization assays, single snapshots of all channels were acquired. For real-time colocalization assays images in 561 nm channel and 488 nm channel were acquired every 5 s with acquisition of the DNA-channel (647 nm) every 1 min for 45 min total (541 frames) using Nikon Elements acquisition software. Single endpoint images were analyzed using ComDet v.0.5.5 plugin for ImageJ (https://github.com/UU-cellbiology/ComDet) to determine colocalization and quantification between DNA channel (647 nm) and GFP (488 nm) and mCherry (561 nm) channels. Results were quantified and plotted using MATLAB (The Mathworks, Natick, MA). Adjustments to example images (contrast, false color, etc.) were made using FIJI [53] and applied equally across entire field of view of each image.

For real-time colocalization assays, a custom-built image analysis pipeline was built in MATLAB (R2019b) to extract DNA-bound intensity traces for the different fluorescent species, to convert them into ON/OFF pulses and to generate the empirical survivor function data [22]. Briefly, the image dataset was drift-corrected using either fast Fourier Transform cross-correlations or translation affine transformation depending on the severity of the drift. DNA spots were identified after binarizing the DNA signal using global background value as threshold, as well as size and time-persistency filtering. Mean values of z-normalized fluorescent markers intensities were measured at each DNA spot at each time frame, and local background was subtracted. Z-normalized traces were then binarized to ON/OFF pulses by applying a channel-specific, manually adjusted threshold value unique to all traces in a given image set. Pulse onsets, durations and overlaps between channels were then derived. For plotting clarity, z-normalized traces shown in the figures were zero-adjusted so that the baseline signal lies around zero. Pulses in ON state at the end of the image acquisition (right censored) were annotated and included as unweighted in lifetime estimates. Kaplan-Meier analysis and log rank tests were performed in MATLAB (R2023a). Adjustments to example plot images (contrast) as well as generation of example plot source movies were made using FIJI [53].

### Live cell microscopy and image analysis

Fixed cell images were acquired on a DeltaVision Ultra deconvolution high-resolution microscope (GE Healthcare) equipped with a 100×/1.42 PlanApo N oil immersion objective (Olympus) and a 16-bit sCMOS detector. Cells were imaged in Z-stacks through the entire cell using 0.2-µm steps using plane-polarized light, 488nm, and 568nm illumination. All images were deconvolved using standard settings. Z projections of the maximum signal in all channels were exported as TIFFs for analysis by in FIJI [53]. 568nm images of Ndc10-mCherry were used to identify centromeric region puncta in cells. The signal intensity within these regions was quantified for the 568nm channel and then for the corresponding 488nm channel images on a per cell basis. Total intensity in the 488nm channels was normalized to total 568nm intensity on a per cell basis and averaged between ∼20cells per biological replicate (total of three biological replicate per strain imaged). Representative images displayed from these experiments are projections of the maximum pixel intensity across all Z images. Adjustments to example cell images (contrast) as were made using FIJI [53].

### Fiber-Seq analysis of *Saccharomyces cerevisiae* genomic DNA

In-house Hia5 preparation and Fiber-seq were performed as described [36], with some adaptation for yeast cultures. Briefly, 10mL of yeast cells (WT, *okp1-AID*, and *cse4-AID*) were grown in YPD media to mid-log phase and were treated with 500 μM indole-3-acetic acid (IAA, dissolved in DMSO) for two hours before being harvested by centrifugation. Cells were washed once with cold dH2O and resuspended in cold KPO4/Sorbitol buffer (1M Sorbitol, 50mM Potassium phosphate pH 7.5, 5mM EDTA pH 8.0) supplemented with 0.167% β-Mercaptoethanol. Cells were spheroplasted by addition of 0.15 ug/mL final concentration of Zymolyase T100 (Amsbio) and incubated at 23 °C for 15 min on a roller drum. Spheroplasts were pelleted at 1200 rpm for 8 min at 4 °C, washed twice with cold 1M Sorbitol, and resuspended in 58 uL of Buffer A (1M Sorbitol, 15mM Tris-HCl pH 8.0, 15mM NaCl, 60mM KCl, 1mM EDTA pH 8.0, 0.5mM EGTA pH 8.0, 0.5mM Spermidine, 0.075% IGEPAL CA-630). Spheroplasts were treated with 1uL of Hia5 MTase (200U) and 1.5uL of 32mM S-adenosylmethionine (NEB) for 10 min at 25 °C. Reaction was stopped by addition of 3uL of 20% SDS (1% final concentration) and high molecular weight DNA was purified using the Promega Wizard® HMW DNA extraction kit (A2920). HMW Fiber-seq modified gDNA was sheared using a g-TUBE (520079) and PacBio SMRTbell libraries were then constructed and sequenced as described [57]. Circular consensus sequence reads were generated from raw PacBio subread files and processed as described [58]. Reads were mapped to the April 2011 sacCer3 yeast reference genome. Nucleosomes were then defined using the default parameters of fibertools [37]. To control for minor technical differences in the overall m6A methylation rate between each sample, reads from each sample were subsetted such that the final distribution of per-read methylation rate was identical between each sample (https://github.com/StergachisLab/match-distribution). Fibers overlapping with the center of each sixteen centromeres +/− 500bp were extracted. The nucleosome density was calculated by counting the number of nucleosomes that overlap with each base pair of the region of interest divided by the number of fibers overlapping with that position. The nucleosome density of all sixteen centromeric regions were averaged and plotted in figure 1B.

### Statistical tests

Significance between colocalization percentages in endpoint assays and in-cell normalized fluorescense intensity were determined by two-tailed unpaired t-tests. Right-censored lifetimes were included in an unweighted Kaplan–Meier analysis to estimate survival functions, censored events typically comprised < 5% of observed lifetimes. Survival functions including 95% confidence intervals were plotted using KMplot (Curve Cardillo, 2008) and P-values for comparing the Kaplan– Meier survival plots are provided in the figure legends and were computed using the log-rank test within Logrank (Cardillo, 2008). Different lifetime plots were considered significantly different with a P-value less than 0.05. Significance between off-rates as well as between cellular intensity values were determined by two-tailed unpaired t-tests.

## DATA AVAILABILITY

Custom software written in MATLAB (R2021a) was used for TIRF colocalization residence lifetime analysis and plot generation. The source code is publicly available at https://github.com/FredHutch/Automated-Single-Molecule-Colocalization-Analysis. All primary datasets and associated data used for analysis in all figures are available at Zenodo, https://zenodo.org/ and assigned the identifier(10.5281/zenodo.xxxxxxx). All primary data reported in this paper will be shared by the lead contact upon request.

## ACKNOWLEDGMENTS

All imaging was performed at the Fred Hutchinson Cancer Center Cellular Imaging Core (supported by the Fred Hutch/University of Washington Cancer Consortium P30 CA015704), and we thank Hoku West-Foyle and Lena Schroeder for their experimental help. We thank Shane Neph for his help with the Fiber-seq bioinformatics analysis. We also thank members of the SB and CLA labs for critical reading of the manuscript. ARP was supported by postdoctoral fellowship NIH F32GM136010. CLA was supported by NIH R35GM134842. ABS holds a Career Award for Medical Scientists from the Burroughs Wellcome Fund and is a Pew Biomedical Scholar and was supported by NIH 1DP5OD029630. SB was supported by NIH R35 GM149357 and is also an investigator of the Howard Hughes Medical Institute.

## AUTHOR CONTRIBUTIONS

**Andrew R Popchock**: Conceptualization; resources; data curation; software; formal analysis; validation; investigation; visualization; methodology; writing – original draft; project administration; writing – review and editing. **Sabrine Hedouin**: Conceptualization; data curation; writing – review and editing. **Charles L Asbury**: Conceptualization; resources; methodology; writing – review and editing. **Yizi Mao;** Conceptualization; resources; methodology. **Andrew B Stergachis;** Conceptualization; resources; methodology; writing – review and editing. **Sue Biggins**: Conceptualization; resources; formal analysis; supervision; funding acquisition; investigation; visualization; methodology; writing – original draft; project administration; writing – review and editing

## SUPPLEMENTAL FIGURES

**Supplemental Figure 1.**
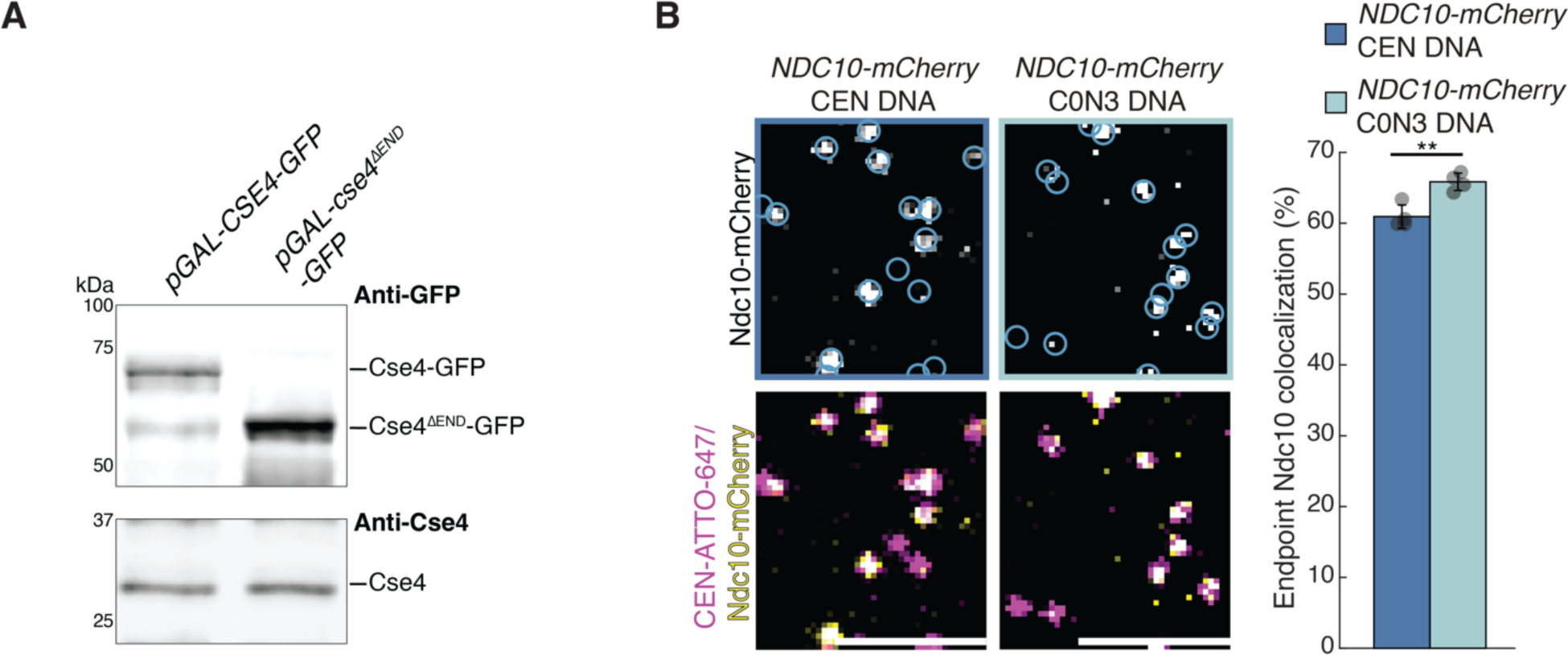
Cse4 recruitment is impaired by loss of the END region or non-native centromeric DNA composition. A. Anti-GFP and anti-Cse4 immunoblots of whole cell extract of *pGAL-CSE4-GFP* (left) and *pGAL-cse4^τιEND^-GFP* (right). B. Example images of TIRFM endpoint colocalization assays. Top panels show visualized Ndc10-mCherry on CEN DNA (top-left panel), or on C0N3 DNA (top-right panel) in *NDC10-mCherry* (SBY8315) extracts with colocalization shown in relation to identified DNAs in blue circles. Bottom panels show overlay of CEN or C0N3 DNA channel (magenta) with Ndc10-mCherry (green), Scale bars 3 μm. Graph indicates quantification of Ndc10-mCherry endpoint colocalization with CEN DNA or C0N3 DNA (61 ± 1.7%, 66 ± 1.2%, avg ± s.d. n = 4 experiments, each examining ∼ 1,000 DNA molecules from different extracts, ** indicates significant difference with two-tailed *P*-value of .003).

**Supplemental Figure 2.**
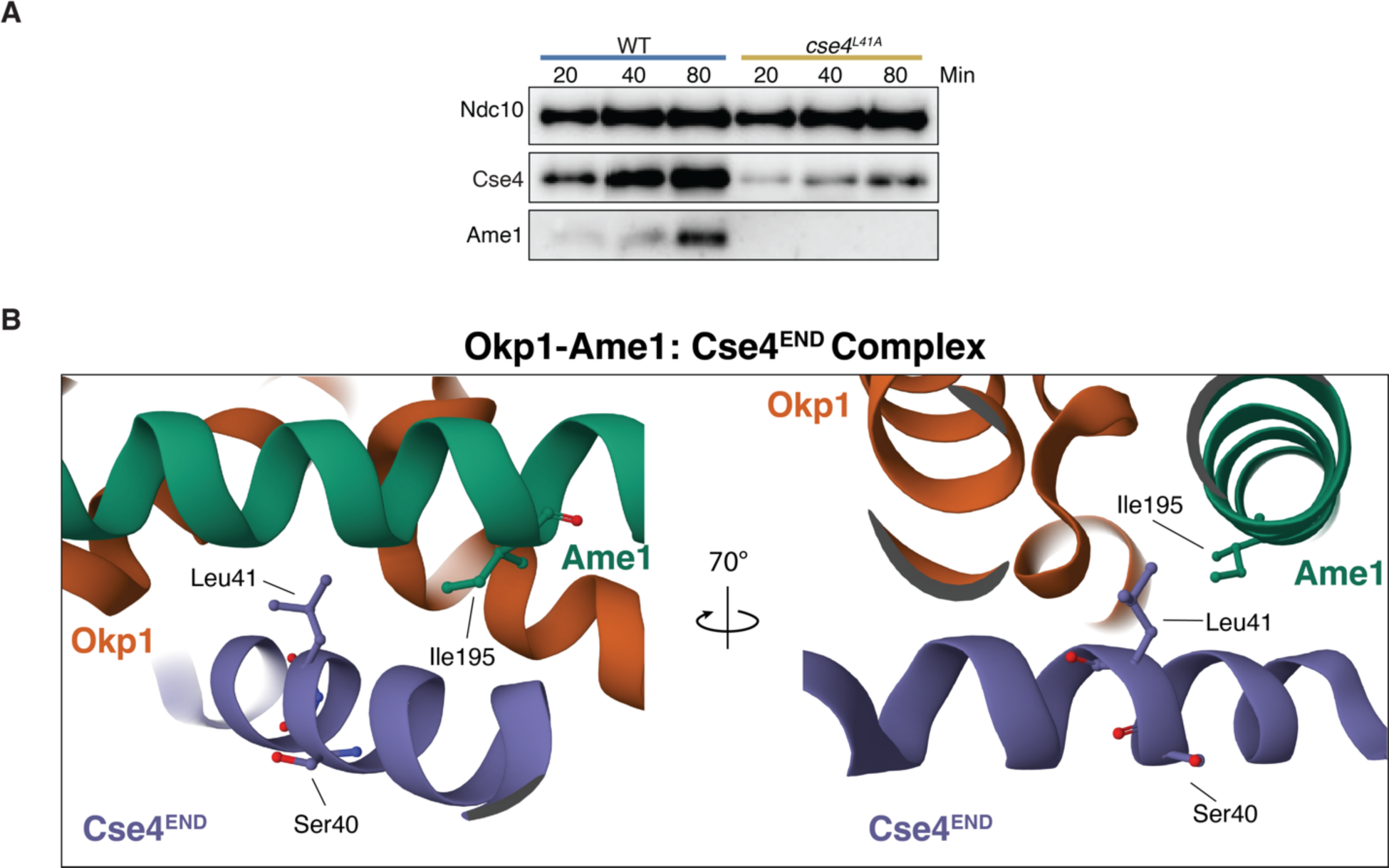
Cse4 and CCAN recruitment is impaired by loss of the END binding to OA during kinetochore assembly. **A.** Immunoblots of bulk kinetochore assembly assays in WT (SBY3) (blue) or *cse4^L41A^* (SBY22466) (orange) extracts on CEN DNA. Centromere DNA-bound proteins were analyzed by immunoblotting with the indicated antibodies. B. Structure of Cse4^END^ (purple) in complex with OA (orange and green), highlighting hydrophobically packed residues Ile195 of Ame1 and Leu41 of Cse4 as well as Cse4 Ser40. Image of 8T0P adapted from [32] and created with Mol* [59].

**Supplemental Figure 3.**
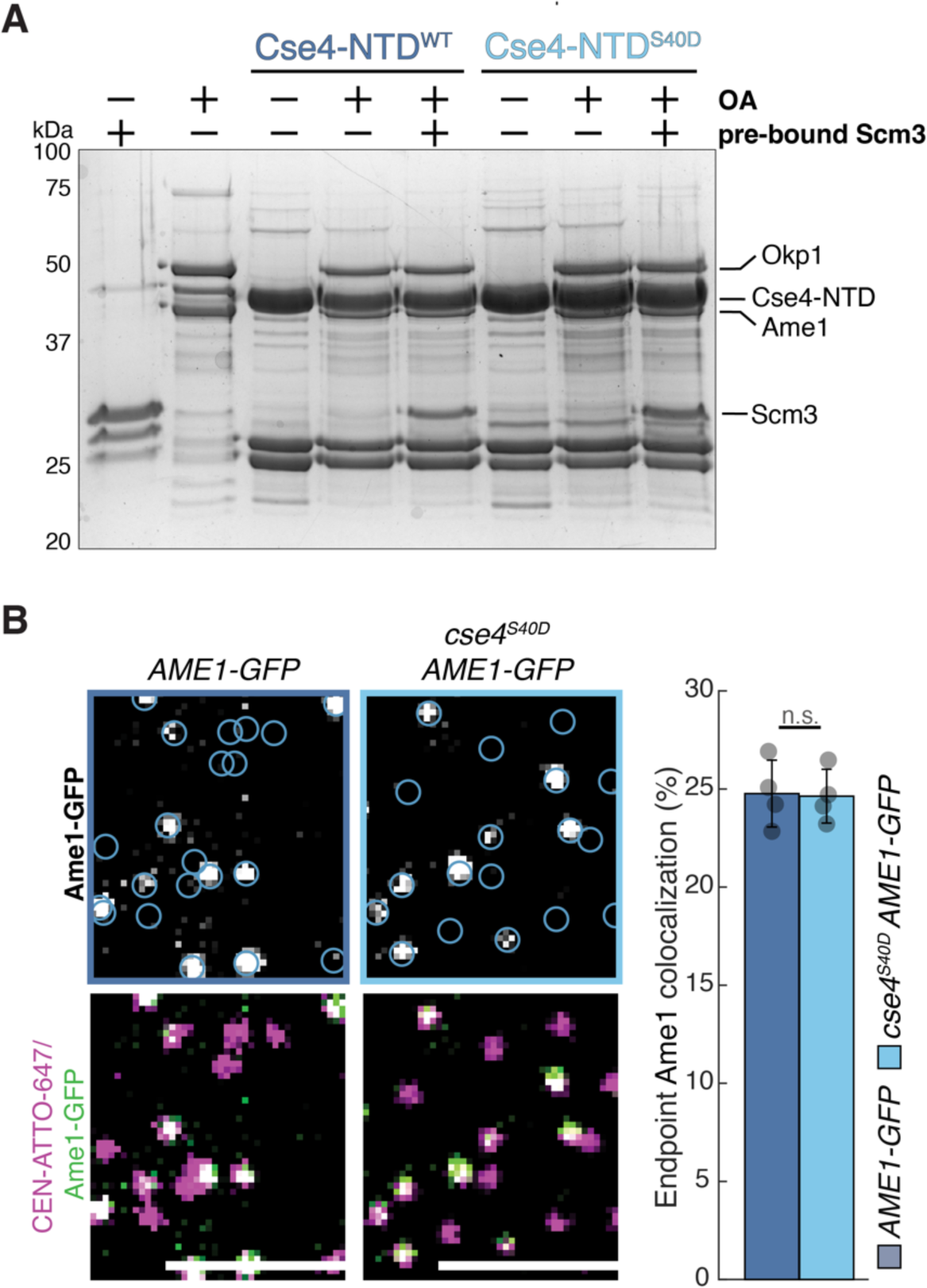
Recruitment of OA is not affected by the Cse4^S40D^ mutant when Scm3 is pre-bound in pulldown assays or in TIRFM assembly assays. A. SDS-PAGE of GST pulldown assays of immobilized Cse4-NTD to test binding of OA with Cse4-NTD^WT^ and Cse4-NTD^S40D^ phosphomimetic mutant alone or in the presence of Scm3. B. Example images of TIRFM endpoint colocalization assays. Top panels show visualized Ame1-GFP on CEN DNA in WT (SBY22273) extracts (top-left panel) or *cse4^S40D^* (SBY22403) extracts (top-right panel) with colocalization shown in relation to identified CEN DNA in blue circles. Bottom panels show overlay of CEN DNA channel (magenta) with Ame1-GFP (green), scale bars 3 μm. Graph indicates quantification of Ame1-GFP endpoint colocalization with CEN DNA in extracts from *AME1-GFP* or *cse4^S40D^ AME1-GFP* genetic backgrounds (25 ± 1.7%, 25 ± 1.4%, avg ± s.d. n = 4 experiments, each examining ∼ 1,000 DNA molecules from different extracts, n.s. indicates two-tailed *P*-value of .9).

**Supplemental Figure 4.**
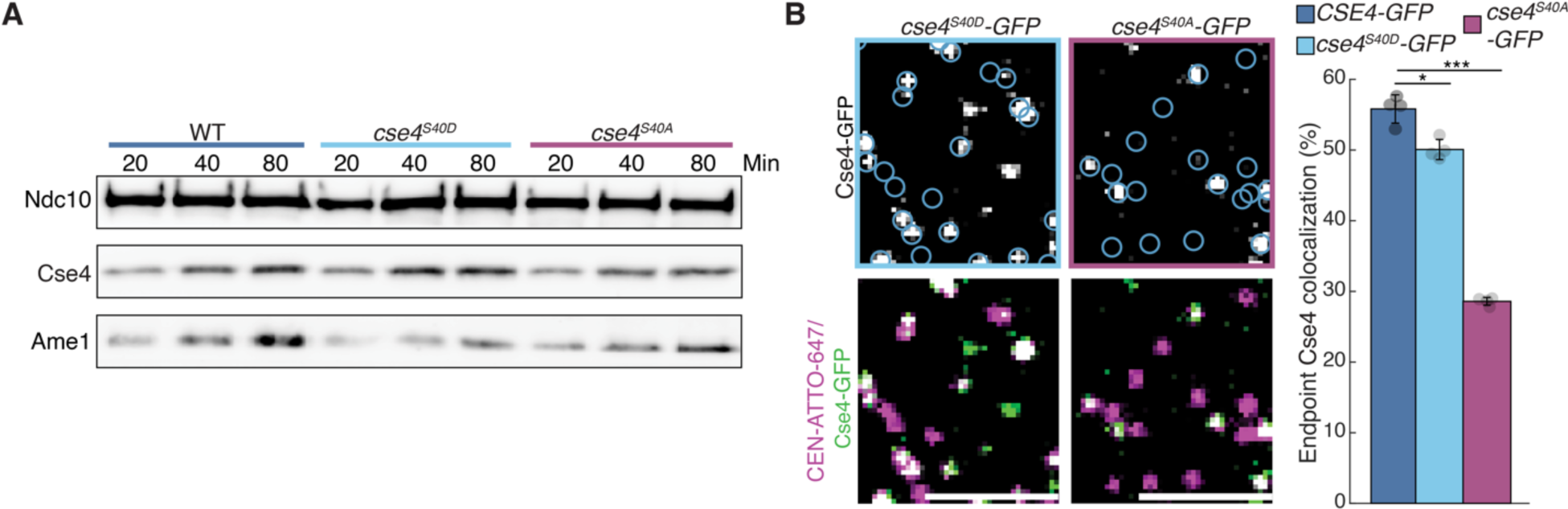
Loss of phosphorylation of Cse4 END at S40 limits centromeric recruitment of Cse4 during kinetochore assembly. A. Bulk kinetochore assembly assays on centromeric DNA in WT (SBY4) (left), *cse4^S40D^* (SBY22401) (middle), or *cse4^S40A^* (SBY22405) (right) cell extracts. DNA-bound proteins were analyzed by immunoblotting with the indicated antibodies. B. Example images of TIRFM endpoint colocalization assays. Top panels show visualized Cse4-GFP in extracts containing *cse4^S40D^-GFP* (SBY20017) (top-left panel) or *cse4^S40A^-GFP* (SBY20019) (top-right panel) on CEN DNA with colocalization shown in relation to identified CEN DNA in blue circles. Bottom panels show overlay of CEN DNA channel (magenta) with Cse4-GFP (green), Scale bars 3 μm. Graph indicates quantification of endpoint colocalization with CEN DNA of Cse4-GFP in extracts containing *cse4^S40D^-GFP* or *cse4^S40A^-GFP* (50 ± 1.4%, 29 ± 0.5% respectively, avg ± s.d. n = 4 experiments, each examining ∼ 1,000 DNA molecules from different extracts, * indicates significant difference between *CSE4-GFP* and *cse4^S40D^-GFP* with two-tailed *P*-value of .006, *** indicates significant difference between *CSE4-GFP* and *cse4^S40A^-GFP* with two-tailed *P*-value of 1.2E-4).

**Supplemental Figure 5.**
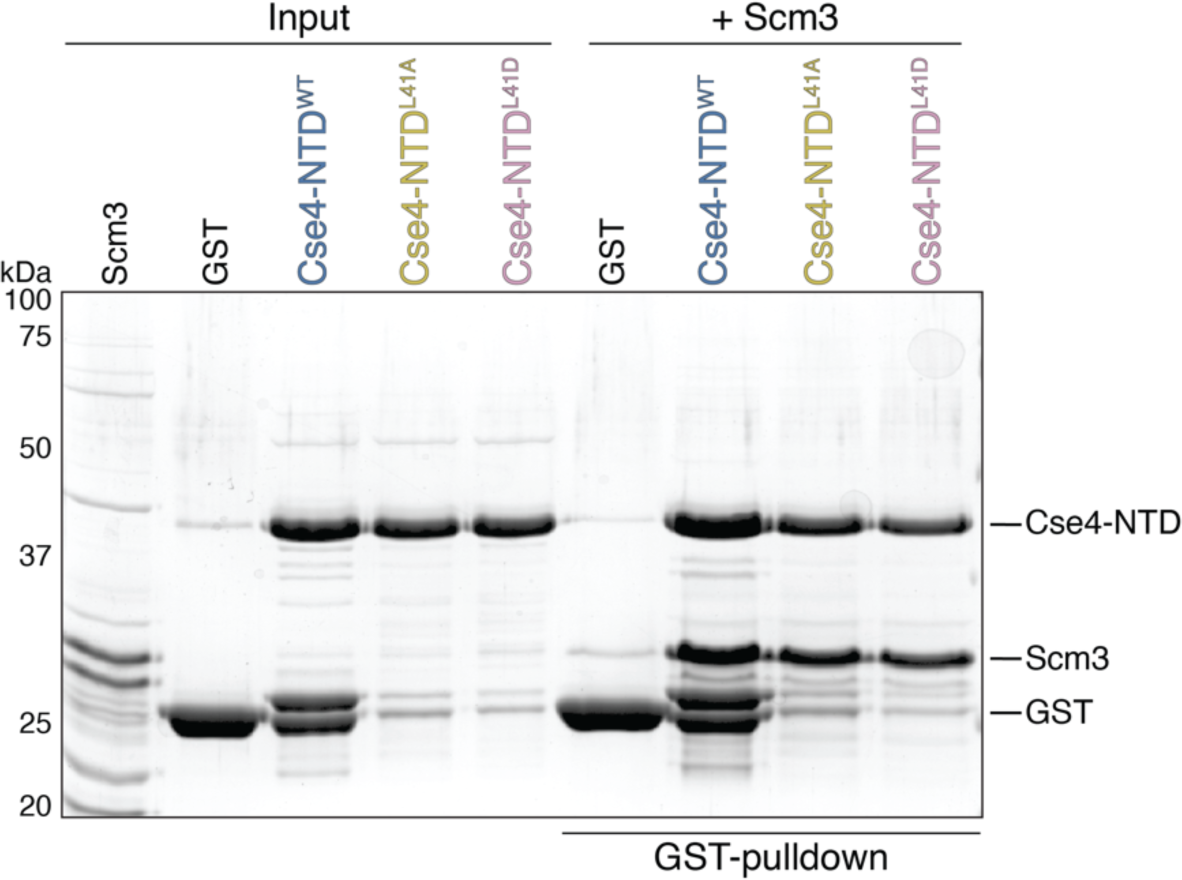
Disruption of Cse4 END binding to OA does not alter Scm3 binding to Cse4-NTD. SDS-PAGE of GST pulldown assays of immobilized Cse4-NTD^WT^, Cse4-NTD^L41A^ and Cse4-NTD^L41D^ to test binding of recombinant Scm3.

**Supplemental Figure 6.**
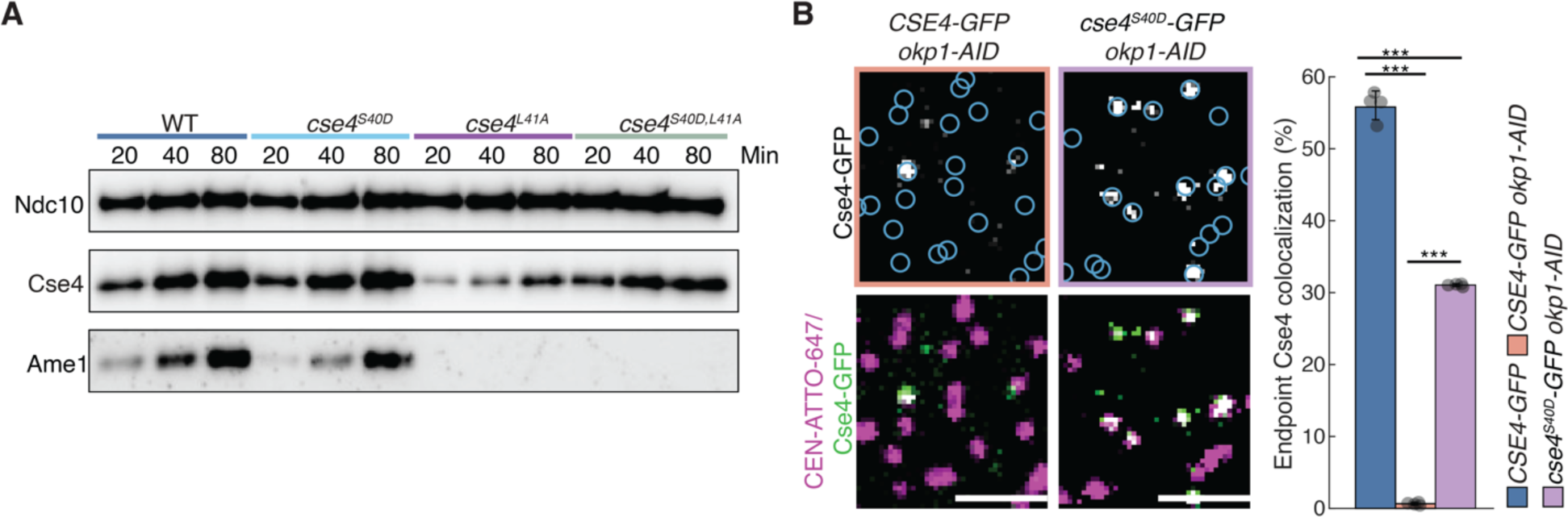
Reduced Cse4 recruitment to centromeric DNA in *cse4^L41A^* mutant or depletion of *OKP1* is rescued by Cse4^S40D^ mutant. A. Bulk kinetochore assembly assays on CEN DNA in WT (SBY21863) extracts, *cse4^S40D^* (SBY20017) extracts, *cse4^L41A^* (SBY22811) extracts, or *cse4^S40D,L41A^* (SBY22914) extracts. DNA-bound proteins were analyzed by immunoblotting with the indicated antibodies. B. Example images of TIRFM endpoint colocalization assays. Top panels show visualized Cse4-GFP on CEN DNA in *CSE4-GFP okp1-AID* extracts (top-left panel), or *cse4^S40D^-GFP okp1-AID* extracts (top-right panel) with colocalization shown in relation to identified CEN DNA in blue circles. Bottom panels show overlay of CEN DNA channel (magenta) with Cse4-GFP (green), scale bars 2 μm. Graph indicates quantification of Cse4-GFP endpoint colocalization with CEN DNA in extracts from *CSE4-GFP*, *CSE4-GFP okp1-AID,* or *cse4^S40D^-GFP okp1-AID* genetic backgrounds (55 ± 0.5%, 1 ± 0.2%, or 31 ± 0.2%, avg ± s.d. n = 4 experiments, each examining ∼ 1,000 DNA molecules from different extracts, *** indicates significant difference between *CSE4-GFP* and *CSE4-GFP okp1-AID* with two-tailed *P*-value of 1.3E-5, between *CSE4-GFP* and *cse4^S40D^-GFP okp1-AID* with two-tailed *P*-value of 1.5E-4, and between *cse4^S40D^-GFP okp1-AID* and *CSE4-GFP okp1-AID* with two-tailed *P*-value of 5.4E-13).

**Supplemental Figure 7.**
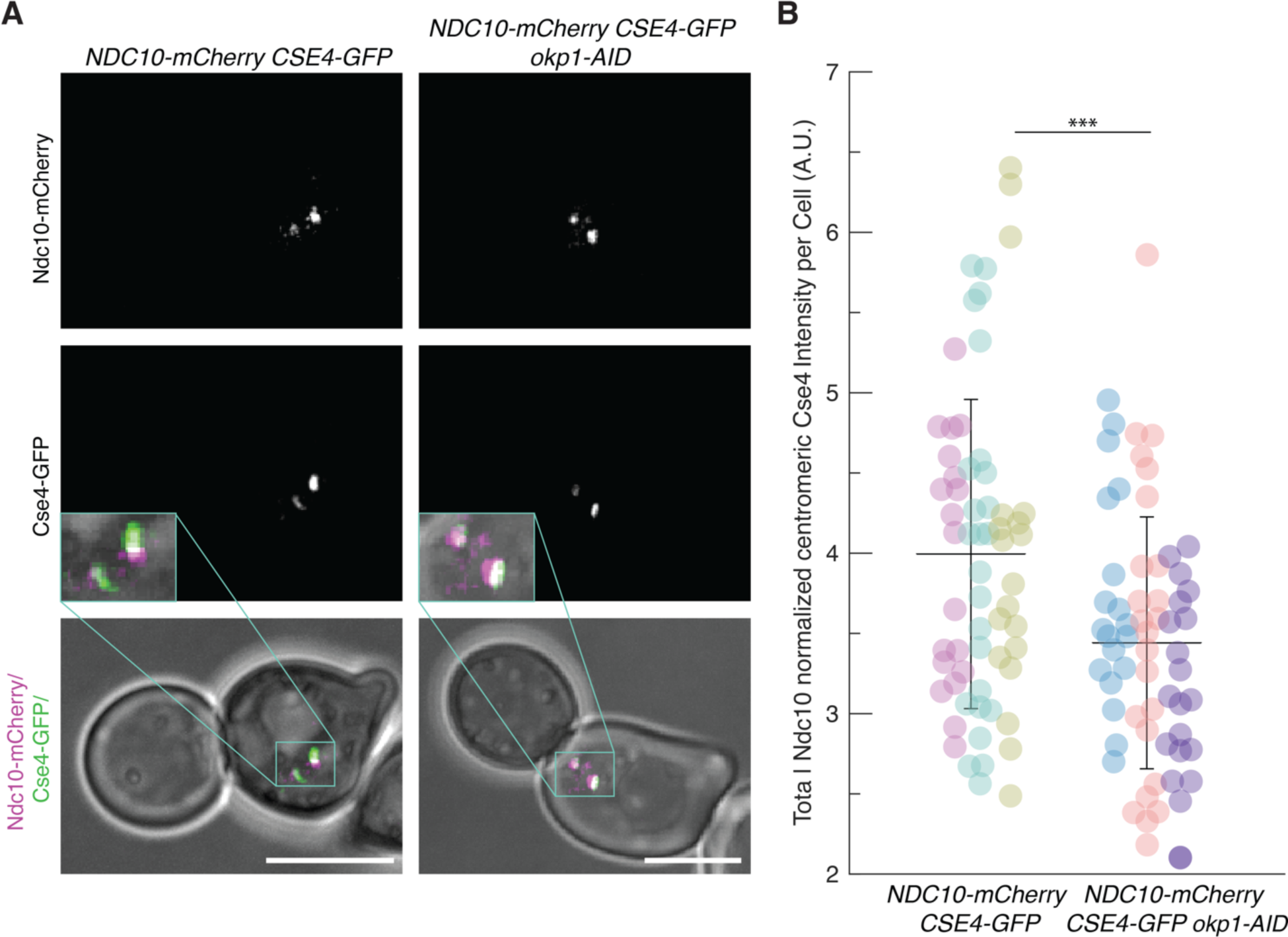
Depletion of Okp1 reduces Cse4 centromeric recruitment in cells. A. Example fluorescence microscopy images of *CSE4*-*GFP NDC10-mCherry* (SBY21974 - left) and *CSE4*-*GFP NDC10-mCherry okp1-AID* (SBY23022 - right) cells showing visualized Ndc10-mCherry (top panels), Cse4-GFP (middle panels) and overlay of Ndc10-mCherry (magenta) and Cse4-GFP (green) on plane polarized illumination of cell. Expanded region around kinetochores highlighted (middle panel inset). Scale bars 5 μm. B. Graph indicates quantification of normalized centromere-associated Cse4-GFP intensity per cell of *CSE4*-*GFP NDC10-mCherry* (left) and *CSE4*-GFP *NDC10-mCherry okp1-AID* (right) cells (4.0 ± 1.0%, 3.4 ± 0.8%, avg ± s.d. n = 3 experiments, each examining ∼ 25 cells). * Indicates significant difference as determined by t-Test (*P*-value of 5.6E-4). Each spot represents calculated intensity for one cell, different colors indicate biological replicates.

**Supplemental Figure 8.**
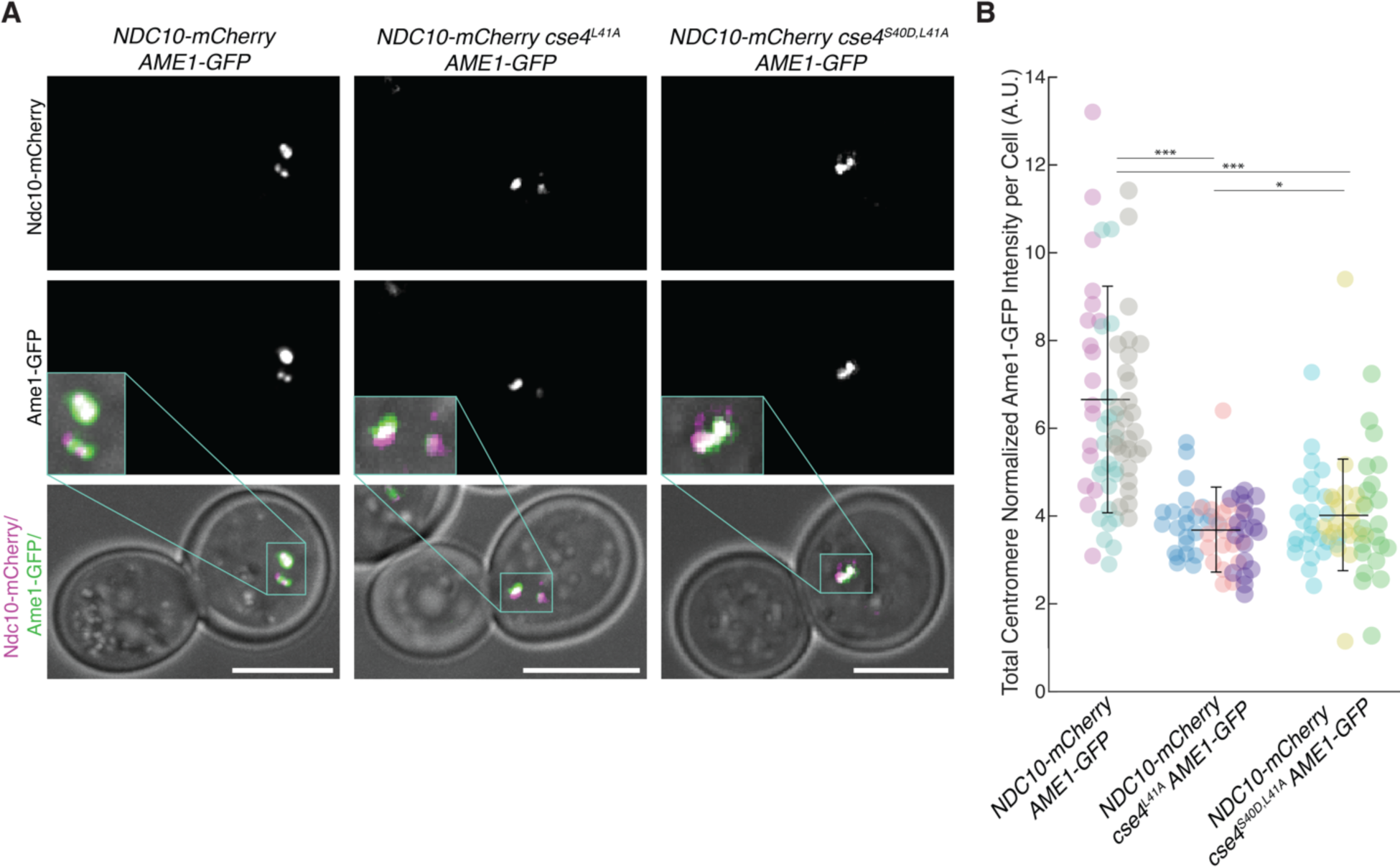
Cse4S40D^,L41A^ mutant does not restore OA localization. A. Example fluorescence microscopy images of *NDC10-mCherry AME1-GFP* (SBY23099 - left), *NDC10-mCherry cse4^L41A^ AME1-GFP* (SBY23295 - middle) and *NDC10-mCherry cse4^S40D,L41A^ AME1-GFP* (right) cells showing visualized Ndc10-mCherry (top panels), Ame1-GFP (middle panels) and overlay of Ndc10-mCherry (magenta) and Ame1-GFP (green) on plane polarized illumination of cell. Expanded region around kinetochores highlighted (middle panel inset). Scale bars 5 μm. B. Graph indicates quantification of Ndc10-mCherry normalized centromere-associated Ame1-GFP intensity per cell of *NDC10-mCherry AME1-GFP* (left), *NDC10-mCherry cse4^L41A^ AME1-GFP* (middle) and *NDC10-mCherry cse4^S40D,L41A^ AME1-GFP* (right) cells (6.7 ± 2.6%, 3.7 ± 1.0%, 4.0 ± 1.3%, avg ± s.d. n = 3 experiments, each examining ∼ 25 cells). * Indicates significant difference as determined by t-Test (*NDC10-mCherry AME1-GFP* : *cse4^L41A^ AME1-GFP P*-value of 2.7E-14, *NDC10-mCherry AME1-GFP* : *NDC10-mCherry cse4^S40D,L41A^ AME1-GFP P*-value of 8.6E-12 and *NDC10-mCherry cse4^L41A^ AME1-GFP* : *cse4^S40D,L41A^ NDC10-mCherry AME1-GFP P*-value of .09). Each spot represents calculated intensity for one cell, different colors indicate biological replicates.

**Supplemental Table S1:**
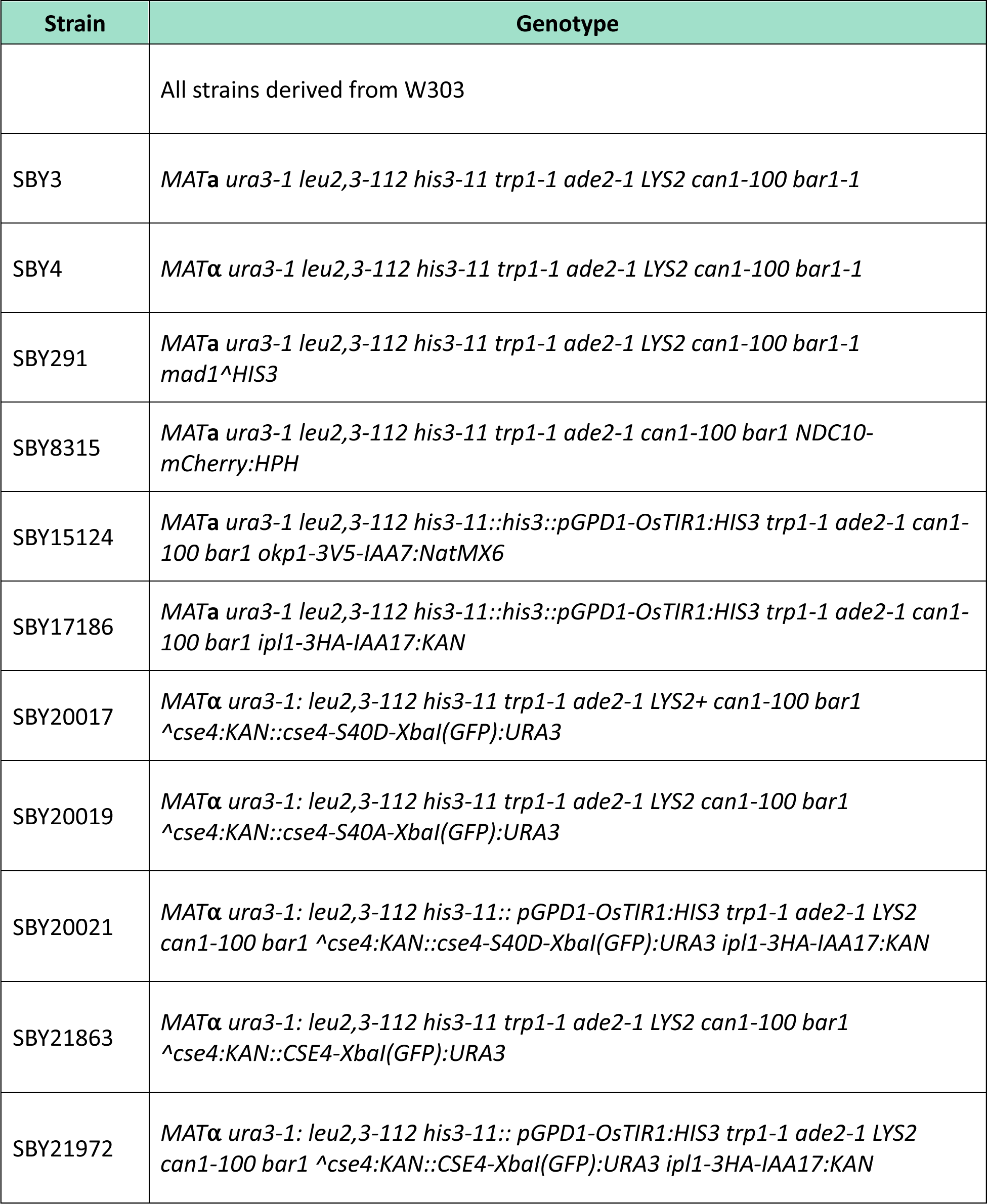
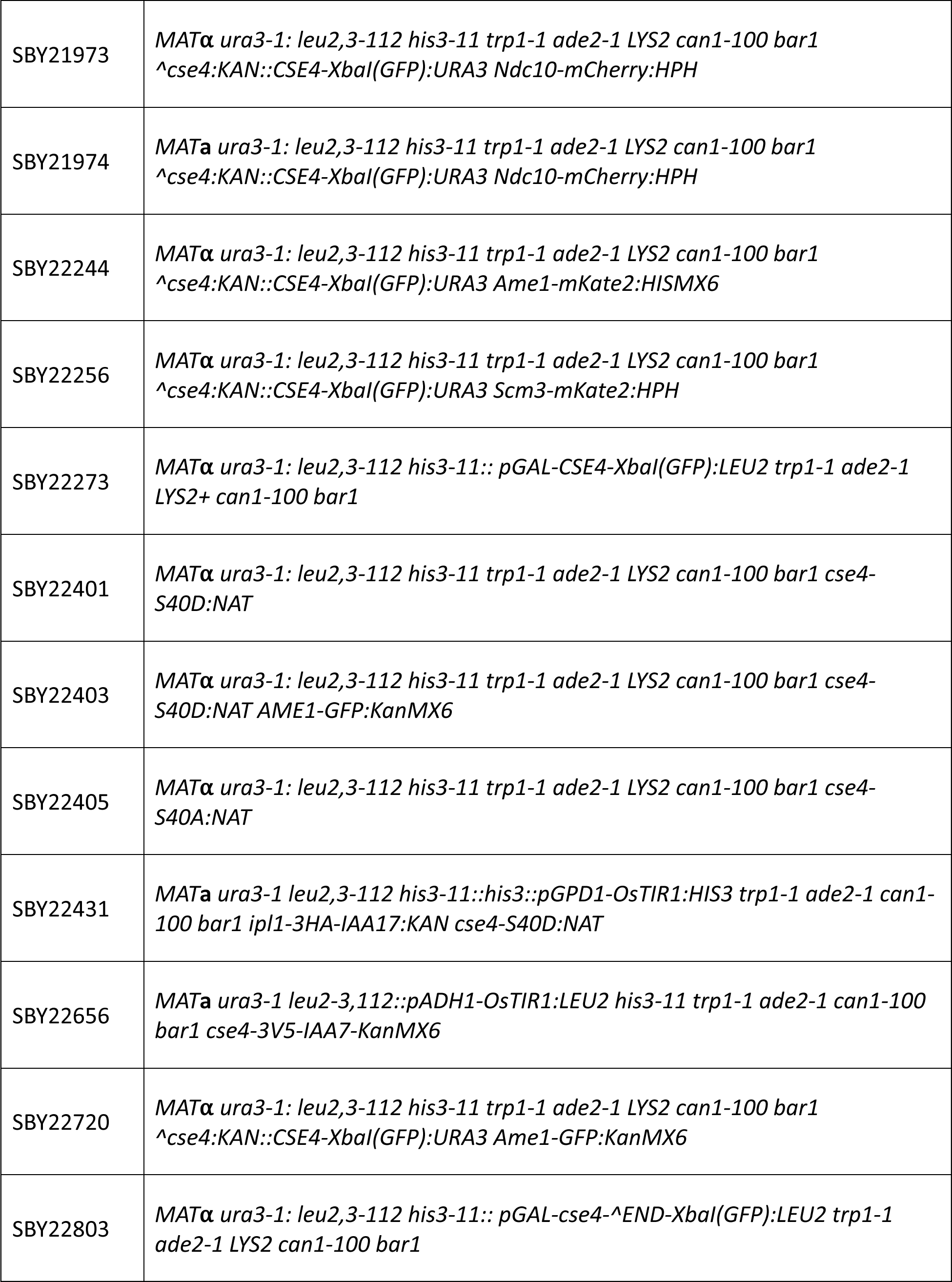
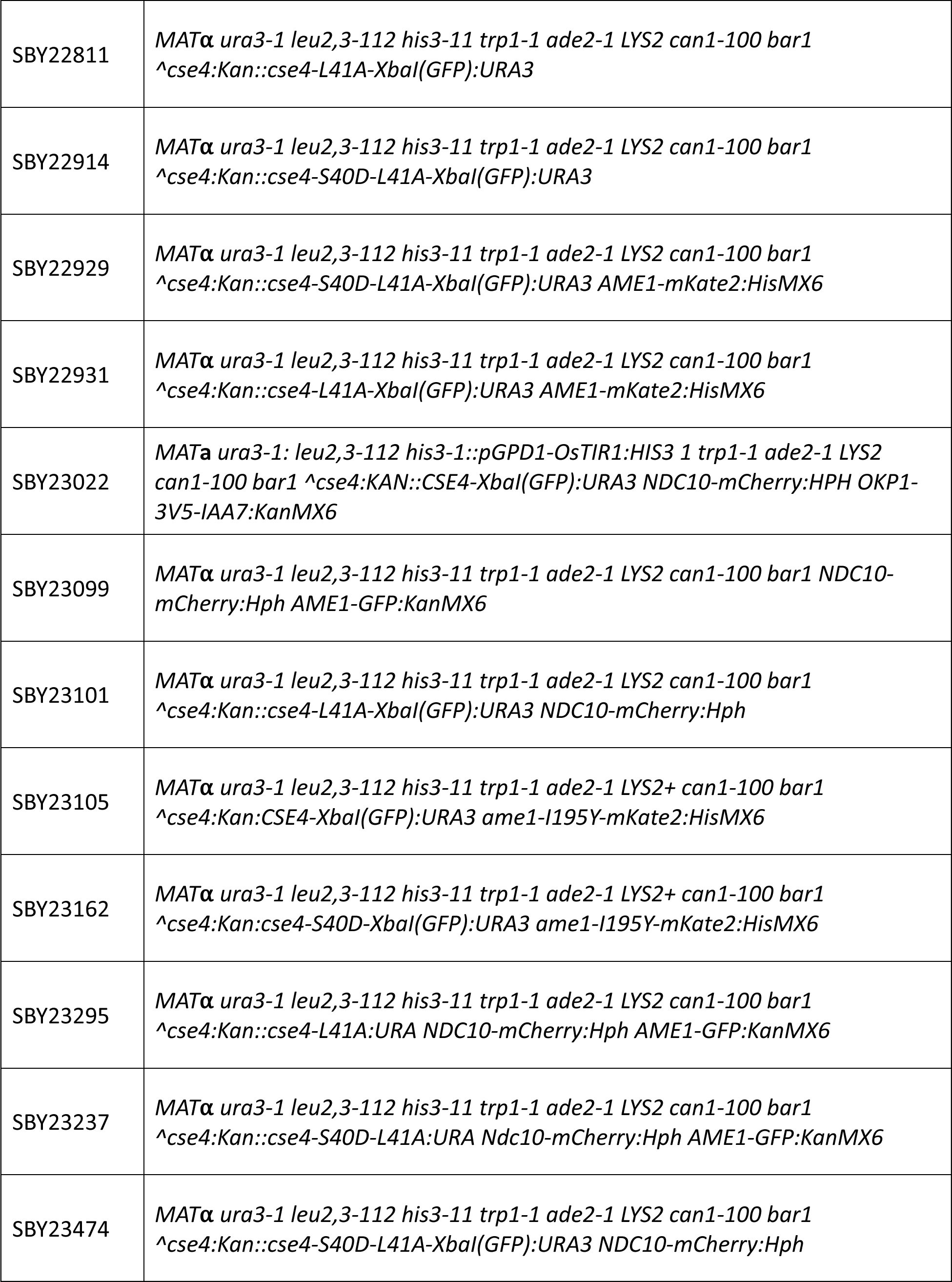
Related to Figures 1-7. List of *S. cerevisiae* strains used in this study.

**Supplemental Table S2:** Related to Figures 1-7. Plasmids used to generate *S. cerevisiae* strains, CEN DNA template generation and recombinant protein expression.

| Plasmid | Description | Source |
| --- | --- | --- |
| pSB64 | *13MYC, HIS3MX6* | Biggins Lab |
| pSB963 | *WT CEN3, 8LacO, TRP1* | Biggins Lab |
| pSB1582 | *mCherry, HPH* | Biggins Lab |
| pSB1617 | *pCSE4-cse4(1-80)-GFP-(81-229), URA3 (integrating)* | Biggins Lab |
| pSB2066 | *3V5-IAA7, KanMX6* | Biggins Lab |
| pSB2273 | *pGPD1-OsTIR1, HIS3 (integrating)* | Biggins Lab |
| pSB3218 | *URA-CEN-gRNA-Cas9 plasmid targeting Cse4* | Biggins Lab |
| pSB3220 | *pCSE4-cse4-S40D-(1-80)-GFP-(81-229), URA3 (integrating)* | Biggins Lab |
| pSB3221 | *pCSE4-cse4-S40A-(1-80)-GFP-(81-229), URA3 (integrating)* | Biggins Lab |
| pSB3249 | *pGAL-cse4(1-80)-GFP-(81-229), LEU2 (integrating)* | Biggins Lab |
| pSB3263 | *pCSE4-cse4(1-80)-mCherry-(81-229), URA3 (integrating)* | Biggins Lab |
| pSB3431 | *mKate2, HISMX6* | Biggins Lab |
| pSB3449 | pET21b - Scm3-6xHIS | Biggins Lab |
| pSB3456 | *pSCM3-SCM3-mKate2 HPH, LEU (integrating)* | Biggins Lab |
| pSB3479 | pET21b - GST | Biggins Lab |
| pSB3480 | pET21b - GST-Cse4(1-131) | Biggins Lab |
| pSB3481 | pET21b - GST-Cse4(1-131)-S40D | Biggins Lab |
| pSB3487 | pET21b - GST-Cse4(1-131)-L41D | Biggins Lab |
| pSB3488 | pET21b - GST-Cse4(1-131)-L41A | Biggins Lab |
| pSB3497 | *pCSE4-cse4-L41A-(1-80)-GFP-(81-229), URA3 (integrating)* | Biggins Lab |
| pSB3498 | pET21b - GST-Cse4(1-27)-(61-131) | Biggins Lab |
| pSB3512 | *EGFP, Kan* | Biggins Lab |
| pSB3526 | *pCSE4-cse4-S40D-L41A-(1-80)-GFP-(81-229), URA3 (integrating)* | Biggins Lab |
| pSB3568 | *pCSE4-cse4-L41A, URA3 (integrating)* | Biggins Lab |
| pSB3569 | *pCSE4-cse4-S40D-L41A, URA3 (integrating)* | Biggins Lab |
| pSB3590 | *pGAL-cse4((1-27)-(61-80))-GFP-(81-229), LEU2 (integrating)* | Biggins Lab |
| pSB3591 | *URA-CEN-gRNA-Cas9 plasmid targeting Ame1* | Biggins Lab |
| pSB3592 | pETDuet - Ame1-6xHIS, Okp1 | Biggins Lab |

**Supplemental Table S3:**
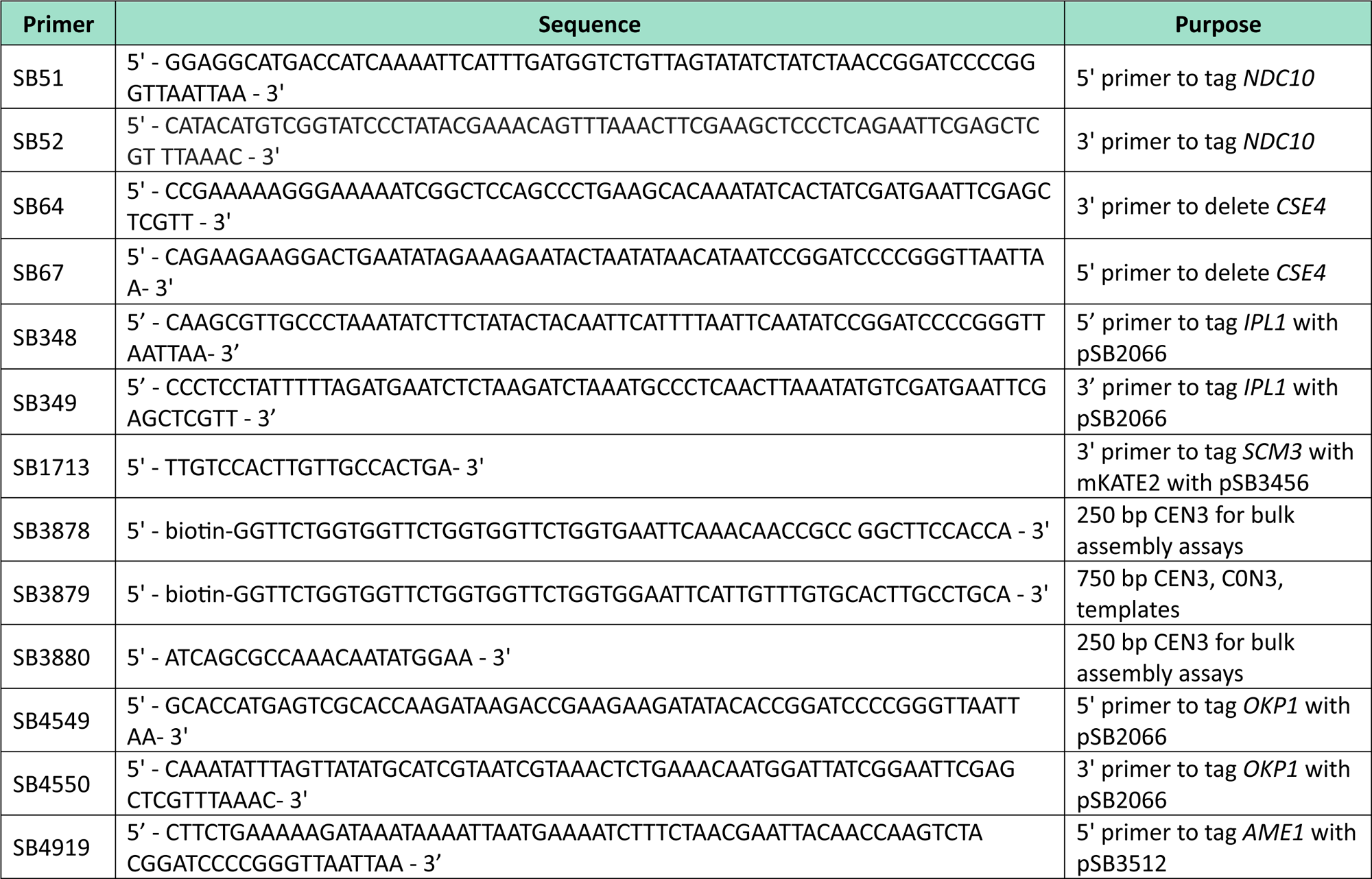
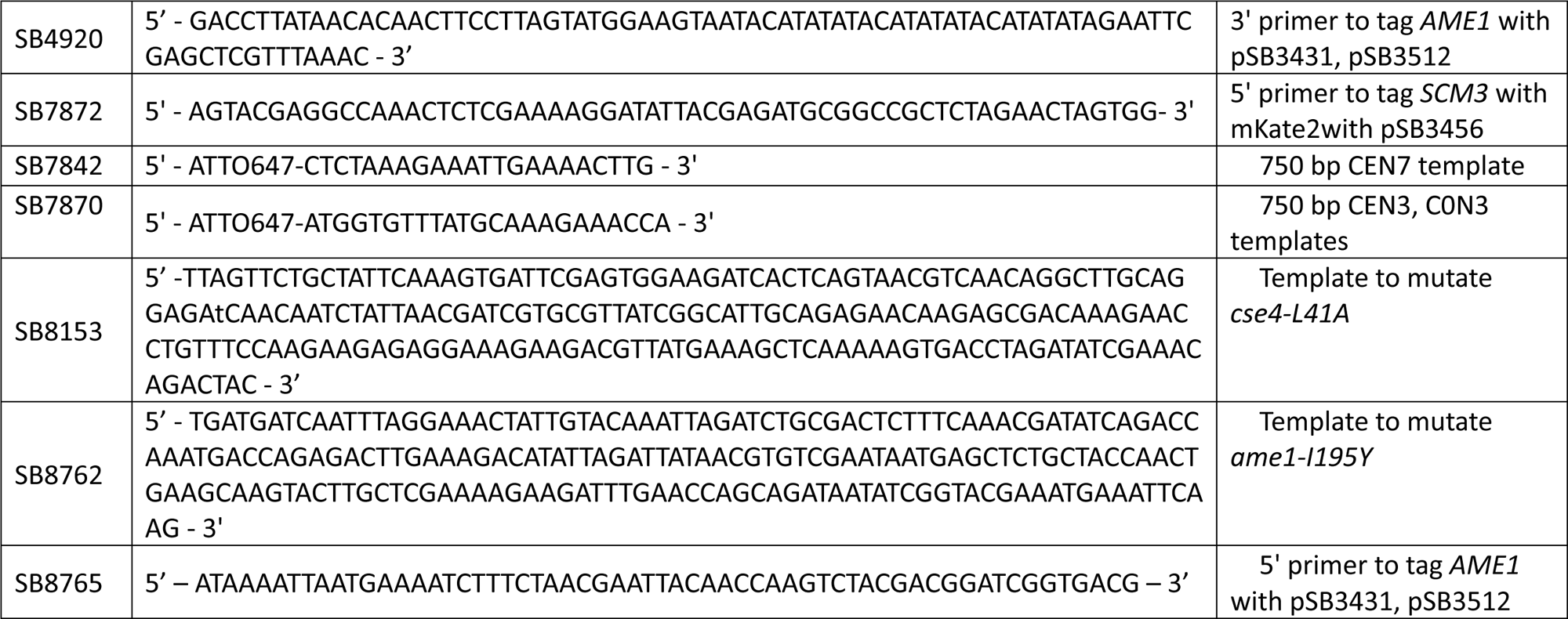
Related to Figures 1-6. DNA oligonucleotides used in this study for *S. cerevisiae* strain construction and CEN DNA template sequence generation.

